# Substrate deformation regulates DRM2-mediated DNA methylation in plants

**DOI:** 10.1101/2020.03.17.995522

**Authors:** Jian Fang, Sarah M. Leichter, Jianjun Jiang, Mahamaya Biswal, Jiuwei Lu, Zhi-Min Zhang, Wendan Ren, Jixian Zhai, Qiang Cui, Xuehua Zhong, Jikui Song

## Abstract

DNA methylation is an important epigenetic mechanism that critically regulates gene expression and genomic stability. In plants, Domains Rearranged Methyltransferase 2 (DRM2) preferentially mediates CHH methylation (H=C, T, A), a substrate specificity distinct from that of mammalian DNA methyltransferases. However, the underlying mechanism is unknown. Here, we report structure-function characterizations of DRM2-mediated methylation. An arginine finger from the catalytic loop intercalates into DNA minor groove, inducing large DNA deformation that impacts the substrate specificity of DRM2. To accommodate the substrate deformation, the target recognition domain of DRM2 embraces the enlarged DNA major groove via shape complementarity, disruption of which via C397R mutation shifts the substrate specificity of DRM2 toward CHG DNA. This study uncovers DNA deformation as a mechanism in regulating the substrate specificity of DRM2, implicative of transposon-specific repression in plants.

## Introduction

DNA methylation at cytosines is an evolutionarily conserved epigenetic mark that critically regulates eukaryotic gene expression and genome stability. Dysregulation of DNA methylation leads to developmental defects and various diseases in animals, most notably cancer, and pleiotropic developmental defects in plants (1-3), highlighting an essential role of DNA methylation in both kingdoms (4, 5). Nevertheless, the mechanisms of DNA methylation have diverged between plants and animals (*5*). In animals, *de novo* DNA methyltransferases DNMT3A and DNMT3B primarily mediate methylation of CpG dinucleotides (6, 7), with appreciable levels of CH (H= A, T, or C) methylation identified in oocytes, embryonic stem cells and neural cells (8). Subsequently, CpG methylation is maintained by DNA Methyltransferase 1 (DNMT1) in a replication-dependent manner (9). In contrast, DNA methylation in plants is prevalent in all sequence contexts: CG, CHG and CHH. Domains Rearranged Methyltransferase 2 (DRM2) mediates the establishment of DNA methylation in all three sequence contexts, whereas plant DNA Methyltransferase 1 (MET1) and Chromomethylase 3 (CMT3) maintain CpG and CHG methylation, respectively. Chromomethylase 2 (CMT2) and DRM2 are jointly responsible for maintaining CHH methylation in long heterochromatic transposable elements (TEs) and short euchromatic TEs, respectively (10). However, the molecular mechanism underlying the divergent methylation patterns between plants and animals remains unclear.

DRM2-mediated methylation is achieved through an RNA-dependent DNA methylation pathway, which involves the biogenesis and enrichment of 24-nucleotide (nt) small interfering RNAs (siRNAs) mostly at short TEs and boundaries of long TEs (11). Targeting DRM2 to specific genomic loci depends on many factors including siRNAs, long noncoding RNAs, histone modifications, and the action of DRM2-interacting proteins (12-15). Emerging evidence has implicated roles of local chromatin environment and the sequence context of DNA substrates in regulating DRM2-mediated methylation. For instance, genome-wide methylation analysis has revealed strong context-dependent methylation in Arabidopsis, with a >900-fold difference between the highest and lowest levels of CHH methylation in the 7-mer sequence context (16). In the example of nucleotide repeat (CCCTAAA)n, the third cytosine has a greater proportion of methylation than the cytosines in the first and second position (16), showing a CHH sub-context specificity. Genomic meta-analysis has also shown that certain trinucleotide contexts, such as CAA and CTA, have a greater methylation frequency than others (e.g. CCC and CCT) (17), supporting a role of sequence context in shaping genomic methylation. Despite these observations, how DNA methyltransferases interplay with substrate sequence to orchestrate distinct DNA methylation patterns at various genomic regions remains unknown.

To elucidate the molecular basis of DRM2-mediated CHH methylation, we performed comprehensive structural characterizations of DRM2-substrate complexes and functional validation analysis *in vivo*. Remarkably, residue R595 from the catalytic core intercalates into the non-target strand, resulting in large DNA deformation, while the target recognition domain (TRD) embraces the deformed DNA major groove via shape complementarity. Biochemical and genome-wide methylation analyses reveal that this DNA deformation permits high methylation efficiency of DRM2 for targets with AT-rich flanking sequences populated in TEs. Substitution of TRD residue C397 with arginine perturbs the shape complementarity between TRD and DNA, shifts the substrate specificity of DRM2 toward the CHG DNA and consequently reshapes the genome-wide DNA methylation patterns. Collectively, this study identified a novel substrate-recognition paradigm for DNA methylation, underpinned by DNA deformation, with strong implication in locus- and sequence-specific DNA methylation establishment and maintenance in plants.

## Results

### Crystal structure of the DRM2-CHH DNA complex reveals substrate deformation

To understand how DRM2 mediates CHH methylation, we determined the crystal structure of DRM2 in complex with a CHH DNA, formed by the methyltransferase (MTase) domain of DRM2 from *Arabidopsis thaliana* (Fig. 1A) and an 18-mer, AT-rich DNA duplex harboring a central <u>C</u>TT motif, in which the cytosine was replaced by a 5-fluorocytosine (5fC) (Fig. 1B). Introduction of the 5fC into the DNA substrate permits the formation of a stable, covalent complex between DRM2 and DNA, as described previously (18, 19). The crystal structure of the DRM2-CTT complex bound to the *S*-adenosyl-homocysteine (SAH), was solved at 2.1 Å resolution (Fig. 1, C and D, and table S1).

**Figure 1.**
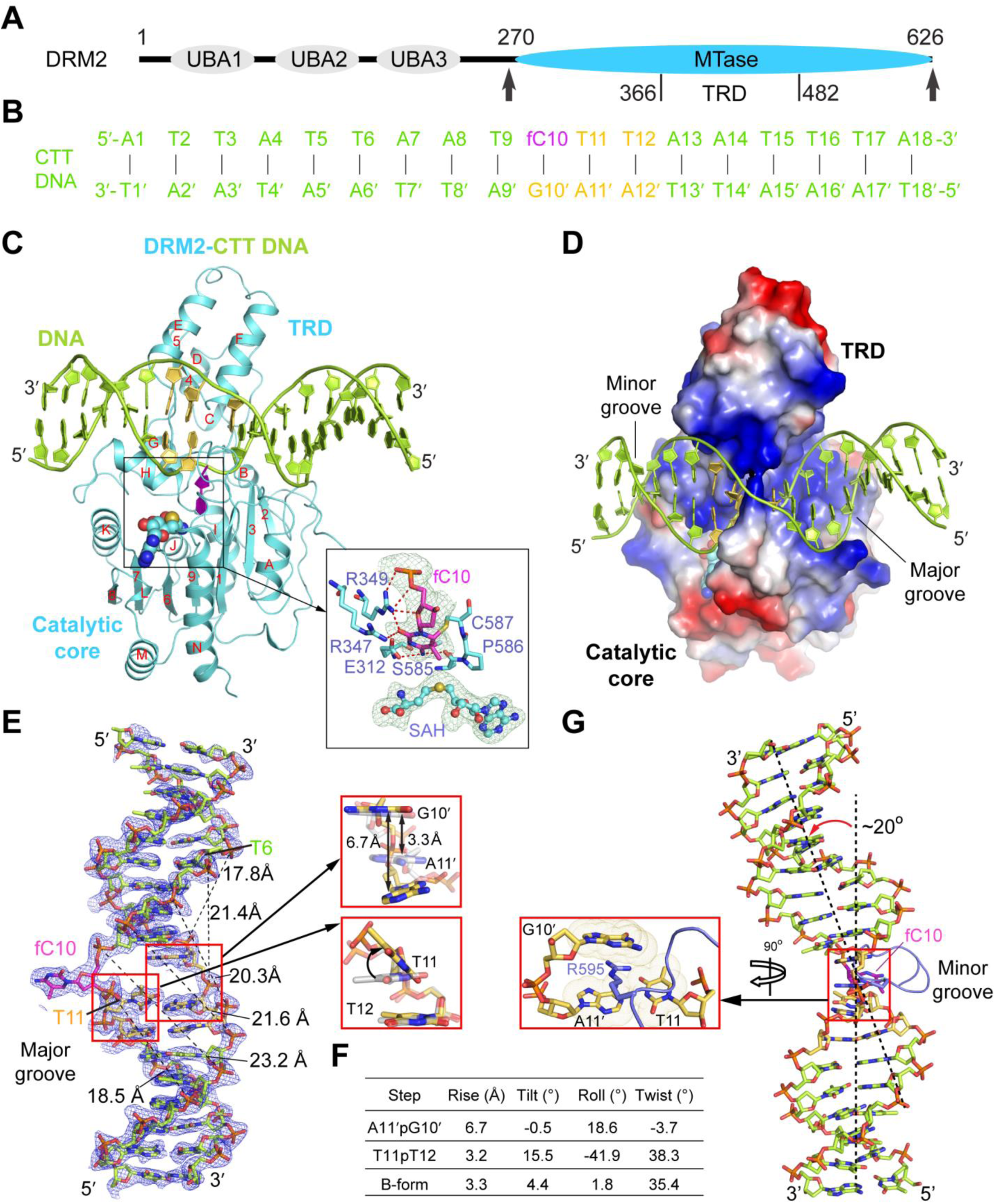
Structure of DRM2 in complex with an 18-mer CTT DNA. (**A**) Domain architecture of DRM2 with the MTase domain marked with arrowheads. (**B**) The sequence of CTT DNA used for the structural study. fC, 5-fluorocytosine. (**C-D**) Ribbon (**C**) and electrostatic surface (**D**) representations of DRM2 bound to DNA and SAH. DRM2 and bound DNA are colored in aquamarine and limon, respectively. The CHH motif colored in yellow or purple (fC10). The SAH molecule is shown in sphere representation. The active site is shown in expanded view, with the Fo-Fc omit map (cyan) of fC10 and SAH contoured at 2.0 σ level and hydrogen-bonding interactions depicted as dashed lines. The color scheme in (C) is applied to subsequent figures, unless otherwise indicated. (**E**) The Fo-Fc omit map (blue) of the CTT DNA, contoured at 2.0 σ level. The major-groove width of the deformed DNA upon binding of DRM2 are measured and marked with dash lines. Structural alignment of the A11′pG10′ and T11pT12 steps with B-form DNA (gray) are shown in expanded views. (**F**) Geometric parameters for the DNA base steps boxed in (E). (**G**) Kinked conformation of DRM2-bound DNA, with the R595 intercalation shown in expanded view.

We were able to trace the entire MTase domain of DRM2 and the DNA molecule. DRM2 is comprised of a catalytic core adopting a Rossmann fold and a target recognition domain (TRD) (Fig. 1C and fig. S1A), as previously observed for DNA-free tobacco DRM (NtDRM) (14). The DNA duplex is embedded in the cleft formed by the TRD and catalytic core of DRM2, resulting in ∼1747 Å^2^ of buried surface area (Fig. 1D). The target 5fC (fC10) is flipped out of the DNA duplex and inserts into the catalytic pocket of DRM2, where it is trapped through covalent linkage with the catalytic cysteine C587 and hydrogen bonding interactions with other catalytic residues (Fig. 1C). Structural comparison of DNA-bound DRM2 with DNA-free NtDRM reveals that their most dramatic structural difference lies in the C587-residing catalytic loop (residues 584-598), which is disordered in DNA-free NtDRM but well defined in DNA-bound DRM2 (fig. S1B), indicating a DNA binding-induced folding. In comparison with B-form DNA, the DRM2-bound DNA undergoes large unwinding (Fig. 1, E to G, and fig. S1C), showing increased inter-strand distances at the segment spanning from Thy6 to Thy11 (Fig. 1E). Most notably, the side chain of R595 on the catalytic loop intercalates into the base step of the non-target strand between the orphan guanine (Gua10′) that was normally paired with fC10, and Ade11′ (Fig. 1G), which increased the helical rise of the Gua10′/Ade11′ step by 3.4 Å and kinked the DNA by ∼20° (Fig. 1, E to G). The R595-mediated DNA intercalation also introduced a large roll and tilt to the +1 fC10-flanking nucleotide, Thy11 (Fig. 1, E and F), which increased the propeller twist of Thy11·Ade11′ pair by ∼23°, leading to a reduced base stacking of the Thy11-Thy12 step (fig. S1C).

### Interplay between DRM2-substrate recognition and DNA shape

The interaction between DRM2 and CTT DNA involves both major and minor groove, spanning thirteen base pairs (Fig. 2, A and B). The minor groove is interrogated by the catalytic loop, and the loop harboring the rearranged motif IV (E312-N313-V314) (20) (rearranged loop: residues 312-320) (Fig. 2B). The major groove is embraced by a loop-helix (αE)-helix (αF) (LHH) motif and helix αH from the TRD, both of which span the widened DNA strands (Fig. 2B). The catalytic loop penetrates into the DNA cavity vacated by base flipping, with residue R595 engaging base-specific interactions with Thy11 and the Gua10′ via direct and water-mediated hydrogen bonds, respectively, in addition to intercalation between Gua10′ and Thy11′ of the non-target strand (Fig. 2C). The unpaired Gua10′ is further stabilized by a direct hydrogen bond between its N7 atom and the backbone carbonyl of G592 (Fig. 2C). On the rearranged loop, residue E312 forms a hydrogen bond with fC10, while residue K319 interacts with the backbone phosphates of non-target strand (T7′ and T8′) through hydrogen-bonding and electrostatic contacts (fig. S1D). Furthermore, the rearranged loop inserts residue L316 into the center of the minor groove to interact with the backbone of both DNA strands through van der Waals contacts (fig. S1D). Additional contacts between the catalytic core and DNA involve the N-terminal loop and β7, which interact with the DNA backbone via hydrogen-bonding interactions (fig. S1E). Toward the major groove, the LHH motif extends along the target strand, then diverts by ∼90° at Thy11 to approach the non-target strand (Fig. 2D), resulting in an ‘L’-shape conformation that complements well with the shape of the deformed DNA (Figs. 1D and 2D). The contacts between the LHH motif and DNA involve hydrogen bonds formed between loop residues (N392, C393 and T396) and the target strand (Ade8, Thy9 and Thy11) (Fig. 2D), and mixed polar and non-polar interactions between the two α-helices (S400, A401, Q402, R406, K433, K434 and W435) and both DNA strands (Fig. 2D). Among these, the sulfhydryl group of DRM2 C397 interacts with the CHH site through van der Waals contacts with the base rings of Thy11 and Thy12 (Fig. 2, D and E). Next to the LHH motif, helix αH spans both DNA strands, with its N-terminal end (S470-T472) engaging hydrogen-bonding interactions with the non-target strand and C-terminal end (G479-S481) engaging van der Waals contacts with fC10 (fig. S1F).

**Figure 2.**
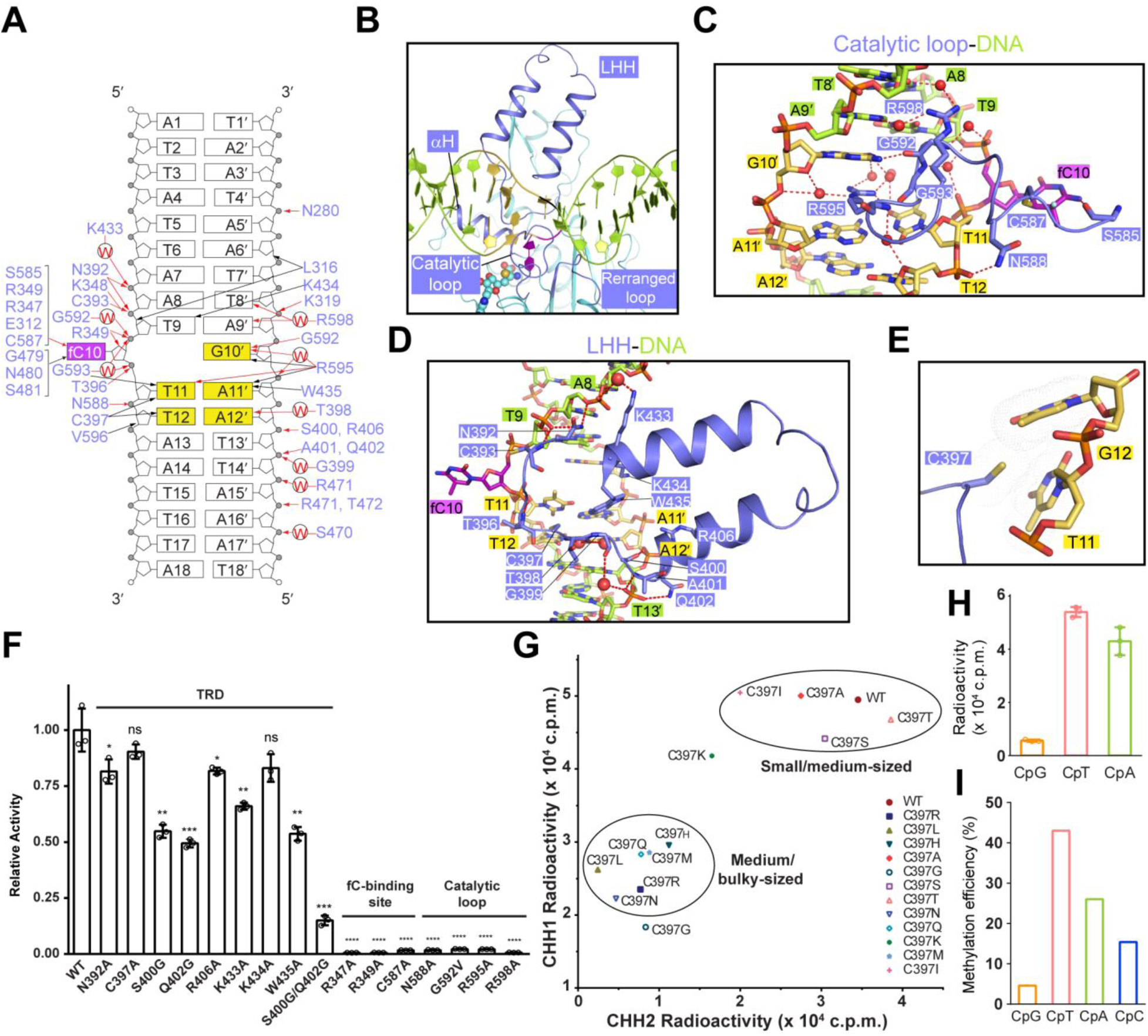
Intermolecular interactions between DRM2 and DNA. (**A**) Schematic view of the intermolecular interactions between DRM2 and CTT DNA. The hydrogen-bonding and van der Waals contacts are represented by red and black arrows, respectively. Water-mediated hydrogen bonds are labelled with the letter ‘W’. (**B**) Close-up view of the DNA-binding regions of DRM2, colored in slate. (**C-D**) Close-up views of the DNA interactions of the catalytic loop (**C**) and LHH (**D**). The hydrogen-bonding interactions are shown as dashed lines. (**E**) Close-up view of the DNA interactions of DRM2 C397. The van der Waals radii of the side chain of C397 and the DNA bases are shown in indicated by dots. (**F**) *In vitro* methylation assay of wild-type (WT) and mutant DRM2. n = 3; Data are mean ± SD. Statistical analysis used two-tailed Student’s t-test for the difference from WT: ns, not significant; *, p < 0.05; **, p < 0.01; ***, p < 0.001; ****, p < 0.0001. (**G**) *In vitro* methylation assay of DRM2, WT or C397 mutants, on two CHH DNA duplexes. CHH1: (TAC)_12_. CHH2: (AAC)_12_. Note that C397K and C397G do not fall into the group classification, likely due to their unique structural, dynamic or charge properties. (**H**) *In vitro* methylation assay of DRM2 on CpG, CpA and CpT DNA. n = 3; Data are mean ± SD. (**I)** Summary of the methylation efficiencies of DRM2 on the CpG/CpH sites of a 637-bp DNA fragment based on bisulfite sequencing analysis. In total 28 clones were analyzed.

To elucidate the interaction between DRM2 and substrates in different sequence contexts, we also solved the structures of DRM2 complexed with <u>C</u>TG, <u>C</u>CG, <u>C</u>AT and <u>C</u>CT DNAs, respectively. The CHG DNAs (<u>C</u>TG and <u>C</u>CG) were derived from the CTT DNA by introducing three or four additional C·G base pairs (fig. S2, A and B), while the new CHH DNAs (<u>C</u>AT and <u>C</u>CT) were derived from the CTT DNA by introducing six or seven additional C·G pairs (fig. S2, C and D). The structures of DRM2-CCT, DRM2-CAT, DRM2-CCG, and DRM2-CTG complexes reveal conserved protein-DNA interactions, and consequently, similar DNA deformation around the CHH/CHG motif (figs. S2-S4, and fig. S5, A to C). Notably, the +2 Gua (Gua12) in the CHG complex is not involved in any protein interaction other than a van der Waals contact with DRM2 C397 (fig. S3, E and F), explaining the lack of specificity of DRM2 toward CHG DNA. Beyond the CHH/CHG motif, the conformation of CTT DNA differs from those of CCG and CTG only at the 5′ flanking region (fig. S5, B and C), but differs from those of CCT and CAT at both flanking regions (fig. S5, B and C), in line with the sequence relationship between these DNAs (fig. S5A). Note that the changes in DNA conformations also lead to altered protein-DNA interactions outside the CHH/CHG motifs (fig. S5, D to F), involving the differential minor-groove contacts at the 5′ flanking region by residues in the N-terminal loop (N280) and rearranged loop (L316 and K319) (fig. S5E), and major-groove contacts at the 3′ flanking region by the LHH (S400, A401, Q402 and R406) and αH (S470-T472) (fig. S5F). Together, these observations not only reveal that DRM2-substrate recognition involves conserved DNA deformation for the CHH/CHG motif, but also point to a role of DNA shape in fine-tuning protein contact by the flanking sequence, an emerging theme in protein-DNA interactions (21).

Next, we mutated key DNA-contacting residues for enzymatic assays. Mutation of the catalytic-loop residues or the fC10-binding site largely abolished the activity of DRM2 (Fig. 2F). Mutation of single TRD residues led to a modest reduction of the enzymatic activity of DRM2, whereas introduction of multi-site mutations (e.g. S400G/Q402G) on TRD severely impaired its activity (Fig. 2F). These data suggest that the catalytic loop plays an essential role in enzymatic catalysis, while the TRD residues collectively stabilize substrate deformation.

The shape complementarity between DRM2 C397 and the +1 - +2 flanking bases (Fig. 2E, and fig. S3, E to H) coincides with the fact that DRM2 proteins from diverse plant species universally contain a small residue (e.g. C, A or V) on the corresponding site (fig. S6). Through mutation of C397 into differently sized amino acids, we observed that the activity of DRM2 on CHH substrates largely falls into two groups, with the smaller amino-acid replacement corresponding to higher activity and larger amino-acid replacement corresponding to lower activity (Fig. 2G). These observations suggest a mechanism in which the shape complimentary between DRM2 and DNA serves to underlie efficient DNA methylation in plants.

### DRM2 methylates DNA in a sequence context-dependent manner

The observation on the DRM2 R595-triggered substrate unwinding is reminiscent of the DNA deformations induced by a group of DNA-binding proteins, such as histones and transcription factors (22, 23). These DNA deformations often occur at A/T-rich regions associated with reduced helical stability (23-25). Coincidently, the Arabidopsis genome is A/T-rich, particularly with small TEs (Fig. S7A), which are the preferential targets of DRM2 (14). Therefore, we ask whether the activity of DRM2 is adapted to such an environment. Toward this, we first interrogated the effect of a +1 flanking base on the enzymatic activity of DRM2 *in vitro*. Among the CpG-, CpT- and CpA-containing DNA, the activity of DRM2 on CpG DNA is much lower than the CT- or CA-containing DNA (Fig. 2H), consistent with a previous observation that DRM2 prefers methylation of CHH and CHG sites over CG sites (26). Furthermore, we measured the activity of DRM2 on a 637-bp DNA containing multiple CG, CC, CT and CA sites via bisulfite sequencing. Our results indicated that DRM2 is most efficient on AT-rich regions (fig. S7B), with the preference for the +1 flanking site following an order of T/A > C/G (Fig. 2I and fig. S7C). These data suggest that DRM2 favors an A/T over a C/G as +1 flanking nucleotide, in line with the general trend for the sequence dependence of DNA unwinding.

To explore the interplay between DRM2 and its target sites *in vivo*, we introduced DRM2 into *drm1drm2cmt3* (*ddc*) triple knockout mutant to determine DRM2-mediated cytosine methylation and the associated flanking sequences. Our whole genome bisulfite sequencing identified 6,792 CG, 16,087 CHG, and 76,234 CHH hypermethylated cytosines (DMCs) mediated by DRM2 (tables S2-S3). These data confirm the A/T-rich environment of the DRM2 targets, with A/T nucleotides accounting for the majority (∼75%) of total nucleotides in the +1 position, and to a less extent in the +2 position (fig. S7D). Together, these data suggest a mechanism of DRM2-mediated methylation that is strongly associated with the sequence context of the methylation sites, and the role of DRM2 in maintaining CHH methylation.

### Role of R595 in DRM2-mediated DNA methylation

To examine the role of R595 intercalation in DRM2-mediated DNA methylation, we compared the enzymatic activities of WT and R595-mutated DRM2. The R595G and R595A mutations largely abolished the activities of DRM2 in all sequence contexts (Fig. 3, A to C). The R595K mutation led to >10-fold reduction of the methylation efficiency on CHH and CHG DNA, but a ∼4.0-fold reduction on CpG methylation (Fig. 3, A to C), confirming the critical role of the side chain of R595 in the enzymatic activity and specificity of DRM2. We further introduced R595G, R595A or R595K into *ddc* triple knockout mutant, which shows global reduction in CHH and CHG methylation and a curled leaf phenotype due to the reactivation of *SDC* (<u>s</u>uppressor of *drm1 <u>d</u>rm2 <u>c</u>mt3*) (27). *SDC* has 7 tandem-repeats in its promoter and is silent in the wild-type plants, but becomes demethylated and transcriptionally reactivated when both DRM2 and CMT3 pathways are inactivated (27). Compared to the wild-type DRM2 transgene that rescued *ddc*, R595G, R595A and R595K all failed to rescue *ddc’s* curled leaf phenotype (Fig. 3D), despite having similar protein levels (Fig. 3E). Consistently, R595A, R595G and R595K showed greatly elevated *SDC* transcript levels similar to *ddc* (Fig. 3F). We next examined the DNA methylation levels of *SDC* and two other DRM2 targets by McrBC digestion and found similar amplification fragments in R595A, R595G, R595K and *ddc* at all these loci (Fig. 3G), indicating that these regions lack DNA methylation. These data highlight the importance of R595 in DRM2 activity, in line with its strict sequence conservation among plant species (Fig. S6).

**Figure 3.**
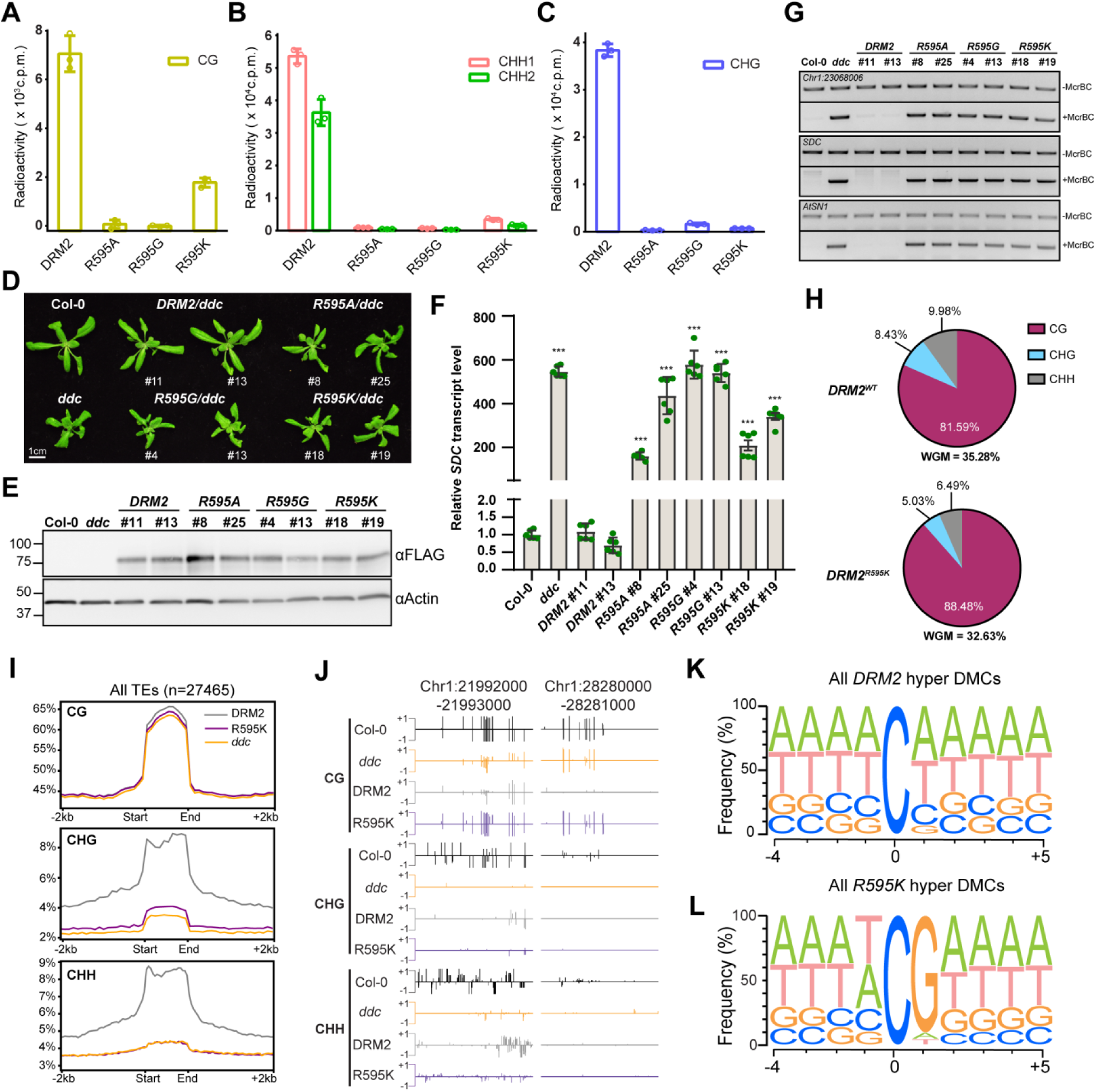
R595 is critical for the activity and specificity of DRM2. (**A-C**) *In vitro* methylation analysis of WT or mutant DRM2 on CG (A), CHH (B) and CHG DNAs (C). CG: (GAC)_12_; CHH1 (TAC)_12_; CHH2: (AAC)_12_; CHG: (TGC)_12_. (**D**) Phenotypes of 3-week-old plants. Col-0 and *ddc* (*drm1drm2cmt3*) serve as controls. (**E**) Western blot of FLAG-tagged DRM2 proteins from same lines listed in panel (D). (**F**) RT-qPCR of *SDC* relative transcript level. Data are mean ± SD. Statistical analysis used two-tailed Student’s t-test for the difference from WT. ***, p < 0.001. (**G**) McrBC digestion of three DRM2 target sites. “-McrBC” represents no enzyme and serves as a control. (**H**) Pie charts showing the proportion of each methylation type in DRM2 and R595K. WGM: whole genome methylation. (**I**) Metaplots showing average methylation level of DRM2, R595K, and *ddc* over all TEs (n=27465) for CG, CHG, and CHH contexts. (**J**) Representative genomic regions showing the methylation levels. (**K**) Motif showing the nucleotide frequency of the 4 nucleotides upstream and 5 nucleotides downstream surrounding hyper differentially methylated cytosines (DMCs) in DRM2 called against *ddc* (n=99103). (**L**) Motif showing the nucleotide frequency of the 4 nucleotides upstream and 5 nucleotides downstream surrounding of hyper DMCs in R595K called against *ddc* (n=1305).

We have further interrogated R595K-mediated genomic methylation through bisulfite sequencing, which revealed a genome-wide methylation decrease in R595K (32.6%) compared to wild-type DRM2 (35.3%) (Fig. 3H). Interestingly, the proportion of methylation context differs greatly between R595K and DRM2, with 88.5% CG in R595K and 81.6% CG in DRM2, accompanied with decreased proportion of CHG and CHH in R595K compared to DRM2 (Fig. 3H). Inspection of the methylation distribution over TEs also revealed a loss of CHH methylation in R595K similarly to *ddc* (Fig. 3I and fig. S8A). On the other hand, R595K exhibited lower reduction of CG and CHG methylation compared to *ddc* (Fig. 3, I and J, and fig. S8A). In fact, flanking sequence analysis of DMCs mediated by DRM2 indicates that R595K mutation greatly affects the nucleotide composition in the +1 position: compared with DRM2-mediated DMCs that favors A/T over C/G as the +1 flanking nucleotide (Fig. 3K and fig. S7D), R595K-mediated DMCs show G as the most abundant nucleotide in the +1 position (Fig. 3L and fig. S8B). Together, these results suggest that R595-mediated DNA deformation governs the activity and specificity of DRM2.

### C397 modulates context-dependent DNA methylation by DRM2

To elucidate how DRM2 diverges from its mammalian counterpart in CHH vs CG methylation, we performed structural comparison of DRM2-CTT with previously reported human DNMT3A-CG DNA and mouse DNMT1-hemimethylated CG DNA complexes (19, 28). The DRM2-CTT complex aligns well with the DNMT3A-CG complex, with a root-mean-square deviation (RMSD) of 1.5 Å over 328 aligned Cα atoms (fig. S9A). Nevertheless, the two complexes show distinct modes of substrate recognition. Notably, unlike DRM2 that wedges the minor groove open through R595-mediated intercalation, DNMT3A presents smaller V716 for stacking against CpG guanine of the target strand, resulting in much less DNA unwinding around the methylation site (fig. S9A). Furthermore, the TRDs of both DNMT3A and DNMT1 engage hydrogen-bonding interactions for CG-specific recognition (fig. S9, A and B), whereas no base-specific interaction is observed for DRM2 TRD, highlighting divergent substrate-recognition mechanisms between plant and mammalian DNMTs. Intriguingly, structural comparison of DNMT3A-DNA and DRM2-DNA complexes reveals that DRM2 C397 is aligned with DNMT3A R836, one of the CG guanine-interacting residues(28) (fig. S9A). This observation prompts us to ask whether replacement of C397 with an arginine would modulate the enzymatic preference of DRM2 also toward a flanking guanine. To address this, we mutated DRM2 C397 into arginine and performed enzymatic assays. Remarkably, the C397R mutation led to reduced enzymatic efficiency toward CHH DNA, but a substantially increased activity toward CHG DNA (Fig. 4A). This data, while reinforces the notion that DRM2-mediated CHH methylation prefers small amino acids at the position of C397 (Fig. 2G and fig. S6), suggests that the C397R mutation alters the substrate recognition of DRM2.

**Figure 4.**
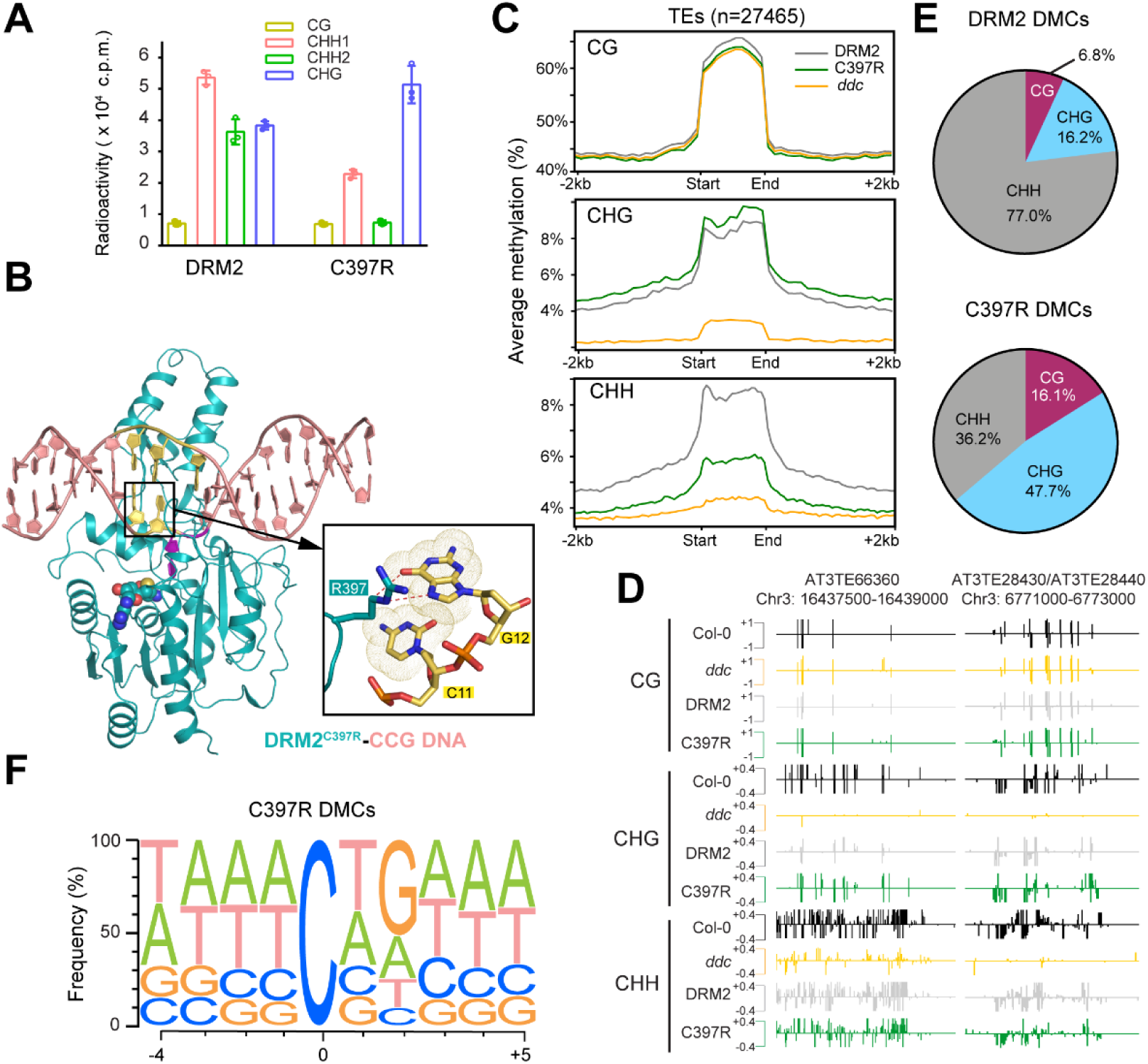
The C397R mutation boosts DRM2-mediated CHG methylation. (**A**) *In vitro* methylation assay of DRM2 or C397R on DNA with different sequence contexts. CG: (GAC)_12_; CHH1: (TAC)_12_; CHH2: (AAC)_12_; CHG: (TGC)_12_. (**B**) Ribbon representations of C397R DRM2 bound to CCG DNA and AdoHcy. Hydrogen-bonding interactions formed between side chain of R397 and G12 are shown in expanded view, depicted as dashed lines. The bases of C11 and G12 in the expanded view are colored in yellow. (**C)** Metaplots showing average methylation level of DRM2, C397R, and *ddc* over TEs for CG, CHG, and CHH contexts. (**D**) Representative genomic region showing the methylation levels. (**E**) Pie charts showing the proportions of differentially methylated cytosines (DMCs) of DRM2 and C397R. (**F**) Motif of the 4 nucleotides upstream and nucleotides downstream of hyper DMCs in C397R called against *ddc* (n = 29347).

To determine the mechanism by which the C397R mutation affects the substrate preference of DRM2, we determined the crystal structure of C397R DRM2 in complex with a CCG DNA (DRM2^C397R^-CCG) at 2.25 Å resolution (Fig. 4B and table S1). The structure of DRM2^C397R^-CCG aligns well with the DRM2-CCG complex, with an RMSD of 0.15 Å over 329 aligned Cα atoms (fig. S10A). Nevertheless, we observed distinct protein interactions involving the +2 guanine (Gua12) between the two complexes: unlike the DRM2-CCG complex where Gua12 only engages van der Waals contact with C397 (Fig. 2E), the DRM2^C397R^–CCG complex involves base-specific hydrogen-bonding interactions between Gua12 and R397: the N7 and O6 atoms of Gua12 are both in close proximity (3.2-3.4 Å) with the side chain of R397, permitting the formation of a hydrogen bond between the N7 atom of Gua12 and the guanidinium group of R397, as well as a C-H-O hydrogen bond between the O6 atom of Gua12 and the Cγ atom of R397 (Fig. 4B). These observations suggest that the C397R mutant recognizes the DNA substrate through a hydrogen-bond directed mechanism, instead of the shape-directed DNA recognition by DRM2, thereby providing an explanation to the change of substrate preference by this mutation.

To verify this change of preference, we compared DRM2- and C397R-mediated DNA methylation *in vivo*. The C397R mutation did not change the total amount of DNA methylation, *SDC* transcript level and leaf phenotype *in vivo* (fig. S10, B to F). However, it increased the relative abundance of CHG and CG methylation from 8.4% and 81.6% to 9.6% and 82.8%, respectively, but reduced the CHH methylation from 10.0% to 7.6% (Fig. S10F). In contrast, mutation of C397 into histidine (C397H) led to a decrease in both the CHH and CHG methylation levels (fig. S10, B to F). In addition, mutation of C397 into alanine (C397A), which naturally occurs at the equivalent position in NtDRM2 (fig. S6), led to a methylation preference similar to DRM2 (fig. S10, B to F, table S3). We further plotted DNA methylation over TEs and found more CHG methylation in C397R than DRM2, C397A or C397H, accompanied with decreased CHH methylation (Fig. 4, C to D, and fig. S10G). We next called hyper DMCs for C397R and DRM2 against *ddc* and found that the proportions of methylation context differ greatly between them, with 47.7% CHG and 36.2% CHH in C397R but 16.2% CHG and 76.9% CHH in DRM2 (Fig. 4E and table S3). Further examination of the flanking sequences around all C397R methylated cytosines revealed a striking difference in the +2 position with 52% being G in C397R (Fig. 4F and fig. S10H) but only 18% G in DRM2 (fig. S7D), supporting the importance of C397-mediated shape complementarity with DNA in substrate discrimination.

Collectively, we have shown structural and functional evidence towards a novel paradigm for substrate recognition of DRM2, with implication for locus-specific DNA methylation in plants.

## METHODS

### Protein expression and purification

A synthetic DNA fragment encoding the MTase domain of *Arabidopsis thaliana* DRM2 (residues 270-626) was cloned into pRSFDuet-1 vector (Novagen), preceded by an N-terminal His_6_-SUMO tag. The expression plasmid was transformed into *Escherichia coli* BL21 DE3 (RIL) cells, and the cells were grown at 37 °C. After the cell density reached an OD_600_ of 0.8, the temperature was lowered to 16 °C. Subsequently, the cells were induced by 100 µM IPTG and continued to grow overnight. The cells were collected and resuspended in Lysis Buffer (50 mM Tris-HCl pH 8.0, 1 M NaCl, 25 mM imidazole, 1 mM PMSF) and lysed using an Avestin Emulsiflex C3 homogenizer. After centrifugation, the supernatant was applied to a Ni^2+^-NTA affinity column and the His6-SUMO-DRM2 fusion protein was eluted with Elution Buffer (20 mM Tris-HCl pH 8.0, 300 mM NaCl, 300 mM imidazole). The His6-SUMO tag was then removed by ubiquitin-like protease 1 (ULP1)-mediated cleavage. The tag-free protein was further purified through ion-exchange chromatography on a Heparin HP column (GE Healthcare) and size exclusion chromatography on a 16/600 Superdex 200 pg column (GE Healthcare). The final protein sample was concentrated and stored in -80°C freezer for future use.

To generate covalent DRM2-DNA complexes, DRM2-MTase, WT or C397R, was reacted with a 18-mer DNA duplex containing a central <u>C</u>TT, <u>C</u>CT, <u>C</u>AT, <u>C</u>CG or <u>C</u>TG motif, in which the target cytosine is replaced by 5-fluorodeoxycytosine (CTT DNA: 5’-ATTATTAATXTTAATTTA-3’; CCT DNA: 5’-ATTCCTCCTXCTCCTTTA-3’; CAT DNA: 5’-ATTCCTCCTXATCCTTTA-3’; CCG DNA: 5’-ATTCCTAATXCGAATTTA-3’; CTG DNA: 5’-ATTCCTAATXTGAATTTA-3’. X = 5-fluorodeoxycytosine), in a buffer containing 25 mM Tris-HCl (pH 8.0), 25% glycerol, 50 mM DTT and 30 µM S-adenosyl-L-methionine (SAM) at room temperature. The reaction products were sequentially purified through a HiTrap Q XL column (GE Healthcare) and a 16/600 Superdex 200 pg column. The final protein samples were concentrated to ∼0.5 mM in a buffer containing 20 mM Tris-HCl (pH 8.0), 250 mM NaCl, 5 mM DTT and 5% glycerol.

### Crystallization conditions and structure determination

For crystallization, the DRM2-DNA complexes were each mixed with 1 mM SAH. Crystals for all of the DRM2-DNA complexes were generated using sitting-drop vapor-diffusion method at 4°C. Each drop was prepared by mixing 0.5 µL of DRM2-DNA complex sample with 0.5 µL of precipitant solution (for DRM2-CTT, DRM2-CTG, DRM2-CCG and DRM2^C397R^-CCG complexes: 2% v/v Tacsimate™(pH 6.0), 0.1 M BIS-TRIS (pH 6.5) and 20% w/v Polyethylene glycol 3,350; for DRM2 ^WT^-CAT complex: 0.1 M Sodium acetate trihydrate (pH 7.0) and 12% w/v Polyethylene glycol 3,350 was used for; for DRM2-CCT complex: 0.1 M Sodium formate pH 7.0, 12% w/v Polyethylene glycol 3,350). The crystal quality was further improved using the micro-seeding method. To harvest crystals, the crystals were soaked in cryoprotectants made of mother liquor supplemented with 30% glycerol, before flash frozen in liquid nitrogen.

X-ray diffraction datasets for the DRM2-CTT, DRM2^C397R^-CCG, DRM2-CCG and DRM2-CCT complexes were collected on beamline 5.0.1 or 5.0.2 at the Advanced Light Source (ALS), Lawrence Berkeley National Laboratory. X-ray diffraction datasets for the DRM2-CAT and DRM2-CTG complexes were collected on the 24-ID-E and 24-ID-C NE-CAT beamlines, respectively, at the Advanced Photon Source, Argonne National Laboratory. The diffraction data were indexed, integrated, and scaled using the HKL 3000 program (29). The structures of the complexes were solved by molecular replacement with the PHASER program (30), using the structure of the MTase domain of NtDRM (PDB 4ONJ) as searching model. The structural models of the DRM2-DNA and DRM2^C397R^-DNA complexes were then subjected to modification using COOT (31) and refinement using the PHENIX software package (32) in an iterative manner. The same R-free test set was used throughout the refinement. The statistics for data collection and structural refinement of the covalent DRM2-DNA and DRM2^C397R^-DNA complexes are summarized in table S1.

### *In vitro* methylation assay

*In vitro* methylation assay was performed in 20 µL reactions containing 1 µM DRM2 (WT or mutants), 3 µM synthesized DNA duplexes containing (GAC)_12_, (TAC)_12_, (AAC)_12_, or (TGC)_12_ sequences to serve as CG, CHH1, CHH2 or CHG substrates, respectively, 0.56 µM *S*-adenosyl-L-[methyl-^3^H]methionine with a specific activity of 18 Ci/mmol (PerkinElmer), 1.96 µM nonradioactive SAM, 50 mM Tris-HCl (pH 8.0), 0.05% β-mercaptoethanol, 5% glycerol and 200 µg/mL BSA. Reactions were incubated at 37°C for 20 min. before quenched by addition of 5 µL of 10 mM nonradioactive SAM to stop each reaction. 12.5 μL of the reaction mixtures were then loaded onto DEAE membrane (PerkinElmer) and air dried. The membrane was washed with 0.2 M ammonium bicarbonate (pH 8.2) three times for 5 min. each, deionized water once for 5 min, and 95% ethanol once for 5 min. After air dried, the membrane was transferred into vials containing 4 mL of scintillation buffer (Fisher) and subjected to tritium scintillation recording by a Beckman LS6500 counter. Each reaction was replicated three times.

### Sanger bisulfite sequencing

A 637-bp DNA substrate, containing multiple target sites (upper strand: 15 CpG, 38 CpA, 26 CpT and 29 CpC sites; lower strand: 15 CpG, 57 CpA, 31 CpT and 26 CpC sites), was derived from a fragment of pGEX-6P-1 vector (nucleotides 302-938) via PCR amplification. The methylation assay was performed *in vitro* in 20 µL reaction mixture containing 0.5 µM DRM2, 0.05 µM DNA substrate, 400 µM SAM (Sigma), 50 mM Tris-HCl (pH 8.0), 0.05% β-mercaptoethanol, 5% glycerol and 200 µg/mL BSA. The reaction was incubated at 37°C for indicated durations, followed by bisulfite conversion using EZ DNA Methylation-Gold™ Kit (Zymo Research). The bisulfite-converted DNA upper and lower strands were subsequently amplified by 2xTaq RED DNA Polymerase Master Mix (APEX) using respective sets of primers (For the upper stand: 5’-TTGAAGAAAAATATGAAGAGGATTTGTATGAG-3’ and 5’-CCCCTCCAACACAACTTCC-3’ were used as forward and reverse primers, respectively; for the lower strand: 5’-ACCCACTCCACTTCTTTTCCAATATC-3’ and 5’-AGGGTGTGAGGTGGGAGAT-3’ were used as forward and reverse primers, respectively). The PCR products were cloned into the pCR™4-TOPO™ Vector (Invitrogen) and subjected to sequencing analysis. In total two biological replicates were assayed.

To generate the Weblogos, 56 clones were analyzed to calculate the average methylation level of CpG, CpA, CpT and CpC sites. Out of total 237 cytosine sites, the top 21 most methylated ones were selected to generate Weblogos using the server (http://weblogo.threeplusone.com/create.cgi) (33), spanning from -7 to +8-flanking sites.

### Calculation of DNA shape parameters

The DNA shape parameters were calculated using the web server (http://web.x3dna.org/) (34), with the structures of DRM2-bound DNA molecules as input.

### Plant materials and growth conditions

All *Arabidopsis thaliana* transgenic lines were derived from ecotype Columbia-0 (Col-0). The triple mutant *drm1 drm2 cmt3* (*ddc*) is a gift from Steven Jacobsen (University of California-Los Angeles). Seeds were sown on ½ MS plates containing 1% sucrose and kept at 4°C for two days before being transferred to long-day conditions (16 hours light/8 hours dark) at 22°C. After ten days of growing on plates, the seedlings were transferred to soil and grown under long-day conditions at 22°C.

### Construction of plasmids and generation of transgenic plants

Genomic DNA sequences of full-length DRM2 with the endogenous 1.3-kb promoter were amplified and point mutations to residues C397 and R595 were made by site-directed mutagenesis using overlapping PCR. Wild-type and mutant DRM2 constructs were further cloned into the pCAMBIA1306 vector with a C-terminal 3xFLAG tag by ClonExpress II One Step Cloning Kit (Vazyme, C112). These constructs were then transformed into *ddc* plants via Agrobacterium-mediated floral dip method (35). Homozygous T_3_ generation plants were used for western blotting and RT–qPCR experiments.

### RNA extraction and RT-qPCR

Total RNA was extracted from 10-day old seedlings grown on ½ MS plates under long-day conditions using PureLink RNA Mini Kit (Thermo Fisher Scientific, 12183025). One microgram of total RNA was reverse transcribed into cDNA with Protoscript II (New England Biolabs, M0368L) followed by qPCR with SYBR Green Master Mix (Bio-Rad, 1725124) using CFX96 Real-Time System 690 (Bio-Rad). Relative *SDC* transcript level to *ACTIN7* was calculated via the ΔCt method.

### Protein extraction and western blotting

Total proteins were extracted from rosette leaves of three-week-old plants using 5% SDS and boiling for 10 min at 95°C before running on SDS-PAGE gel. FLAG-tagged proteins were detected with horseradish peroxidase (HRP)-conjugated anti-FLAG antibody (Sigma Aldrich, A8592-1MG). All western blots were developed using the ECL Plus Western Blotting Detection System (GE Healthcare, RPN2132) and chemiluminescent imaging using an Imagequant LAS 4000 (GE Healthcare).

### McrBC digestion

Genomic DNA was extracted from 100 mg of rosette leaf tissue of three-week-old plants using PureLink Plant Total DNA Purification Kit (Thermo Fisher Scientific, 45-7004). 100 ng of genomic DNA was treated with McrBC enzyme (New England Biolabs, M0272L) at 37°C overnight and then for 10 min at 75°C to deactivate the enzyme. Digested DNA and undigested DNA were amplified using genomic loci-specific primers.

### Bisulfite sequencing library construction and data analysis

Bisulfite treatment and sequencing library construction were conducted as previously described (36). Briefly, genomic DNA was extracted from three-week-old plants using DNeasy Plant Mini Kit (Qiagen 69104) and ∼1 µg of DNA was sheared to 300-400bp using the Covaris S220 (Covaris) using the Covaris SonoLab 7.5. Sheared DNA was used to construct the library using the Illumina TruSeq DNA PCR-Free Low Throughput Library Prep Kit (Illumina, 20015962). After adapter ligation, samples were bisulfite treated using EZ DNA Methylation-Gold Kit (Zymo Research, D5006) and then amplified for 10 cycles using Kapa HiFi HotStart Uracil ReadyMix (Kapa Biosystems, KK 2801) before sequencing on a Hiseq 4000 (Illumina) with 50bp single-end reads. Sequencing reads were trimmed using FASTP (37) and aligned to the *Arabidopsis* TAIR10 genome using bsmap version 2.9 (38) allowing for 8% mismatches, trimming anything with a quality score of 33 or less, and removing any reads with more than five N’s. Methylation at every cytosine was called using bsmap’s methratio.py script processing only unique reads and removing duplicate reads. Differentially methylate cytosines (DMCs) were identified using both MethylKit (39) and bsmap’s methdiff.py script with the following cutoffs for each sequence context: 40% difference for CG, 20% difference for CHG, and 10% difference for CHH. DMCs called by MethylKit and methdiff.py were compared using BEDtools (40) intersectbed and only overlapped DMCs were used for subsequent analysis. Flanking sequence surrounding each DMC was found using BEDtools getfasta and the resulting fastas were compiled into a motif using Weblogo (33). TE metaplots were created by deeptools (41) computeMatrix using bsmap methylation file and a list of all TEs from TAIR10.

### Statistics

The two-tailed Student’s t-tests were performed to compare distributions between different groups. And the p value lower than 0.01 was considered to be statistically significant.

### Accession codes

Coordinates and structure factors for the DRM2 complexes have been deposited in the Protein Data Bank under accession codes xxxx, xxxx, xxxx, xxxx, xxxx and xxxx. The bisulfite-sequencing data has been deposited in NCBI Gene Expression Omnibus under accession number GSE146700.

## Acknowledgments

We would like to thank Drs. Jie Liu and Dean Sanders for assistance of BS-seq analysis. We also thank staff members at the Advanced Light Source (ALS), Lawrence Berkeley National Laboratory and the Advanced Photo Source (APS), Argonne National Laboratory for access to X-ray beamlines, and Northwestern University Sequencing Core Facility for high throughput sequencing. This work was supported by NIH grant (1R35GM119721) to J.S, NIH (1R35GM124806) and NSF CAREER (1552455) to X.Z.. Group of JZ is supported by the Program for Guangdong Introducing Innovative and Entrepreneurial Teams (2016ZT06S172).

## Author Contributions

J.F., S.M.L., J.J., M.B., J.L., Z-M.Z. and W.R. performed the experiments, J.Z. provided technical support, Q.C. performed computational analysis, X.Z. and J.S. conceived and organized the study. J.F. S.M.L., J.J., X.Z. and J.S. prepared the manuscript.

## Supplementary Information

**Figure S1.**
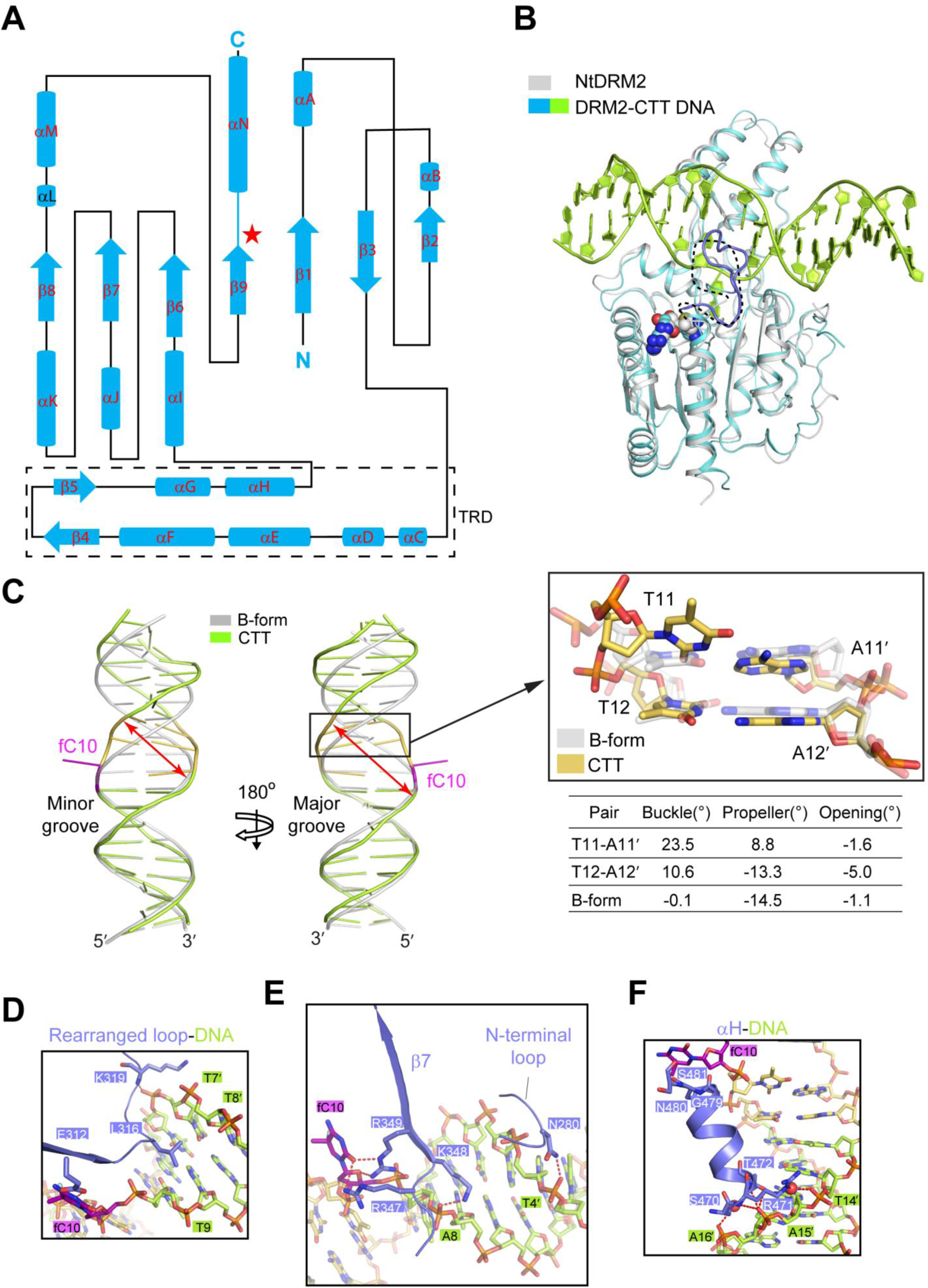
Structural analysis of the DRM2-CTT complex. (**A**) Schematic representation of the secondary structures of DRM2. The active site is marked by red star. The TRD is boxed with dashed lines. The N and C termini of DRM2 are indicated. **(B)** Structural overlay of free NtDRM MTase (PDB 4ONJ) with the 18-mer CTT DNA-bound state of DRM2. The catalytic loop is disordered in free NtDRM MTase but well defined in the DRM2-CTT complex, depicted as dashed lines and loop representation (slate), respectively. The SAH molecule is shown in sphere representation. (**C**) Structural superposition of the DRM2-bound CTT DNA and an Ideal B-form DNA with the corresponding sequence. The enlarged major and minor grooves in the DRM2-bound DNA are indicated by arrows. Detailed structural overlay of the two base pairs next to fC10 between DRM2-bound DNA and B-form DNA is shown in expanded view. The corresponding base-pair parameters are listed below, with those for the ideal B-form DNA listed as reference. (**D-F**) Close-up view of the DNA interactions of rearranged loop (D), β7 and N-terminal loop (E) and αH (F) of DRM2. The hydrogen-bonding interactions are shown as dashed lines. The CHH motif is colored in purple (fC) or yellow.

**Figure S2.**
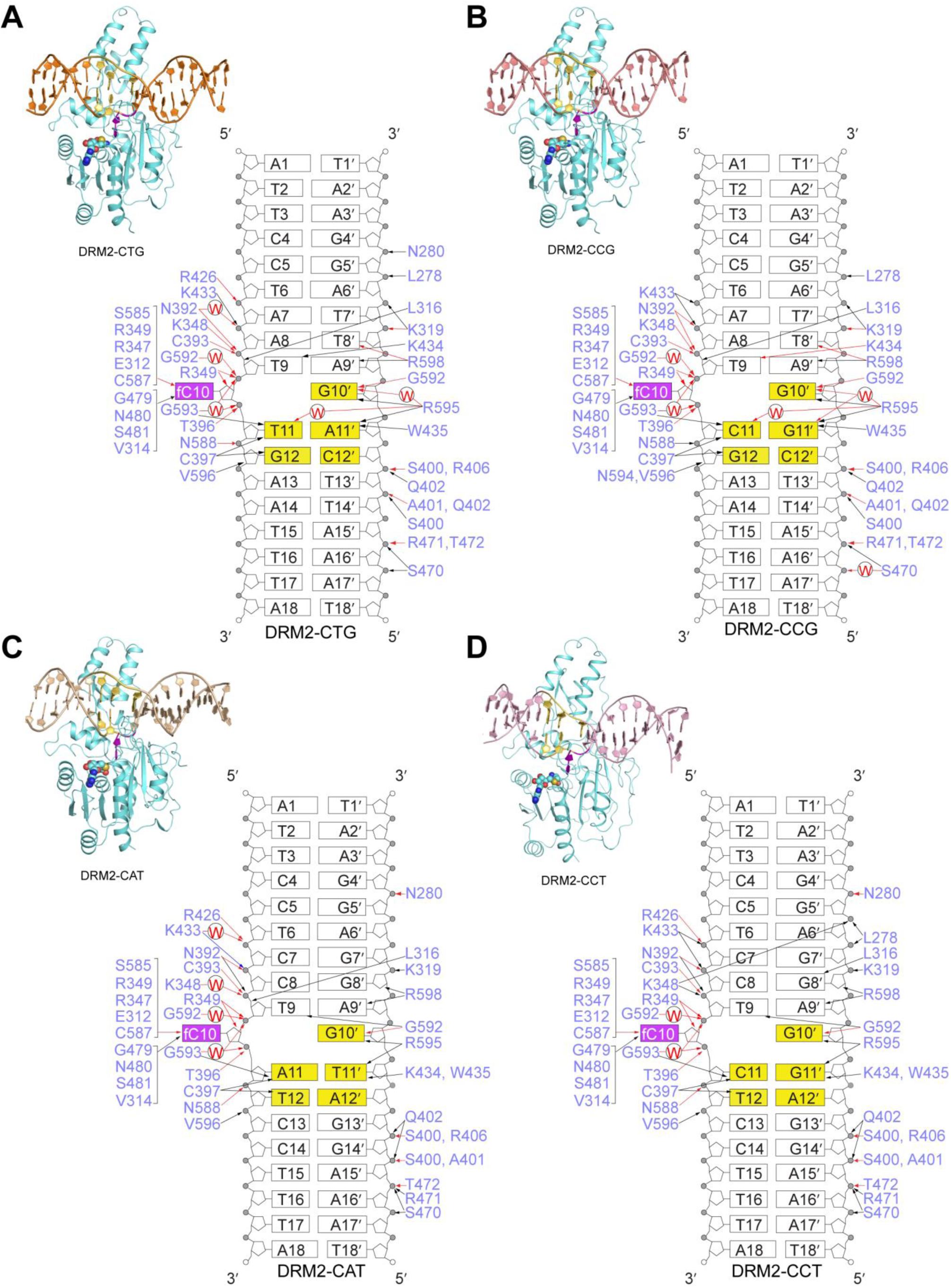
Crystal structures of the DRM2 in complex with CTG, CCG, CAT and CCT DNAs. Ribbon representations of DRM2 in complex with CTG (**A**), CCG (**B**), CAT (**C**) and CCT (**D**) DNA and SAH, with the corresponding DNA sequences and protein-DNA interactions shown in schematic representation. The hydrogen-bonding, electrostatic and van der Waals contacts are indicated by red, blue and black arrows, respectively. Water-mediated hydrogen bonds are labeled with letter ‘W’. The CHH/CHG sites are colored in purple (fC) or yellow.

**Figure S3.**
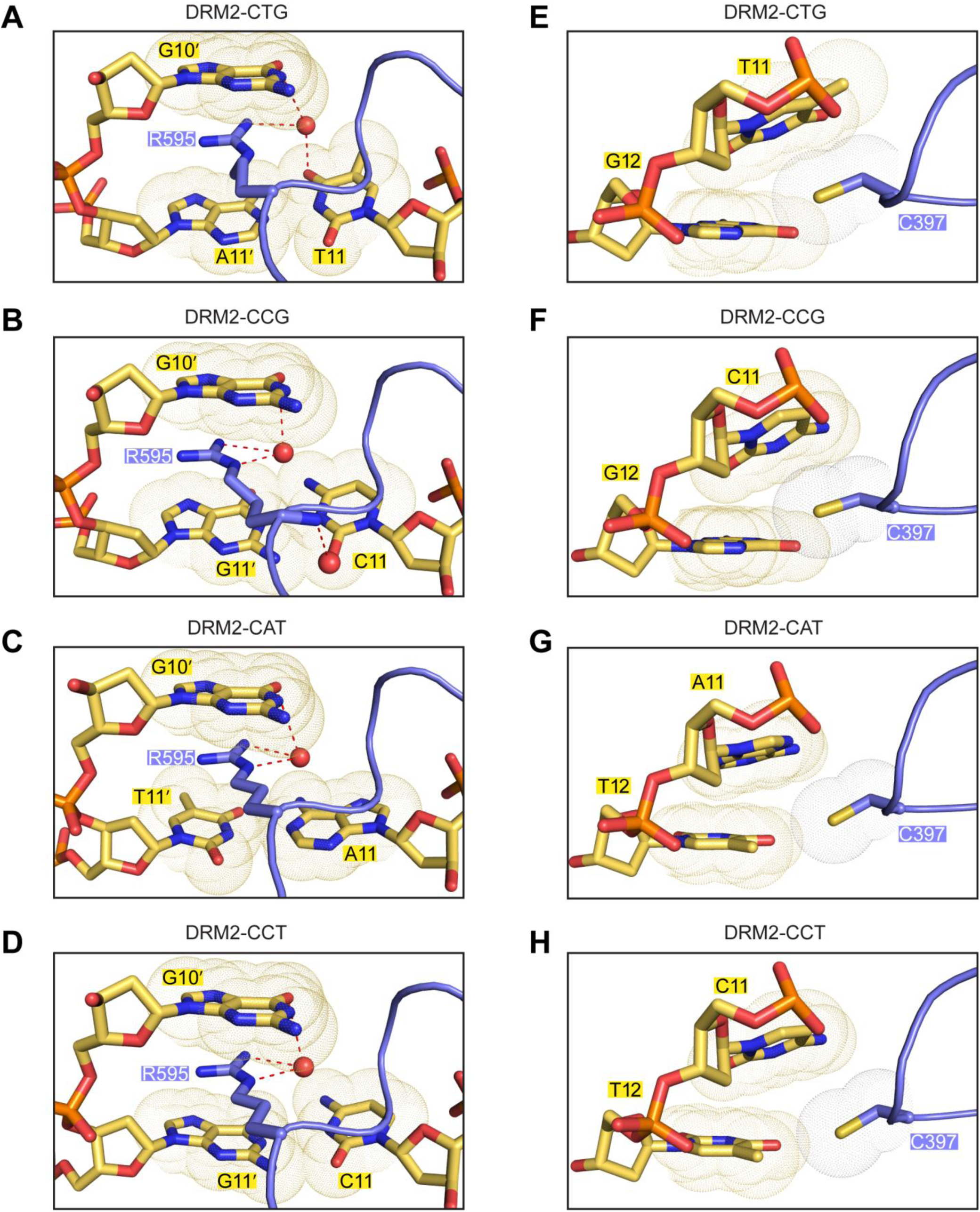
Structural analysis of the R595- and C397-mediated DNA interactions of DRM2. **(A-D)** Close-up views of the DNA interactions of the R595 in the DRM2-CTG (A), DRM2-CCG (B), DRM2-CAT (C) and DRM2-CCT (D) complexes. (**E-H**) Close-up views of the DNA interactions of the C397 in the DRM2-CTG (E), DRM2-CCG (F), DRM2-CAT (G) and DRM2-CCT (H) complexes. The van der Waals radii of DNA bases are indicated by yellow dots. The hydrogen-bonding interactions are shown as dashed

**Figure S4.**
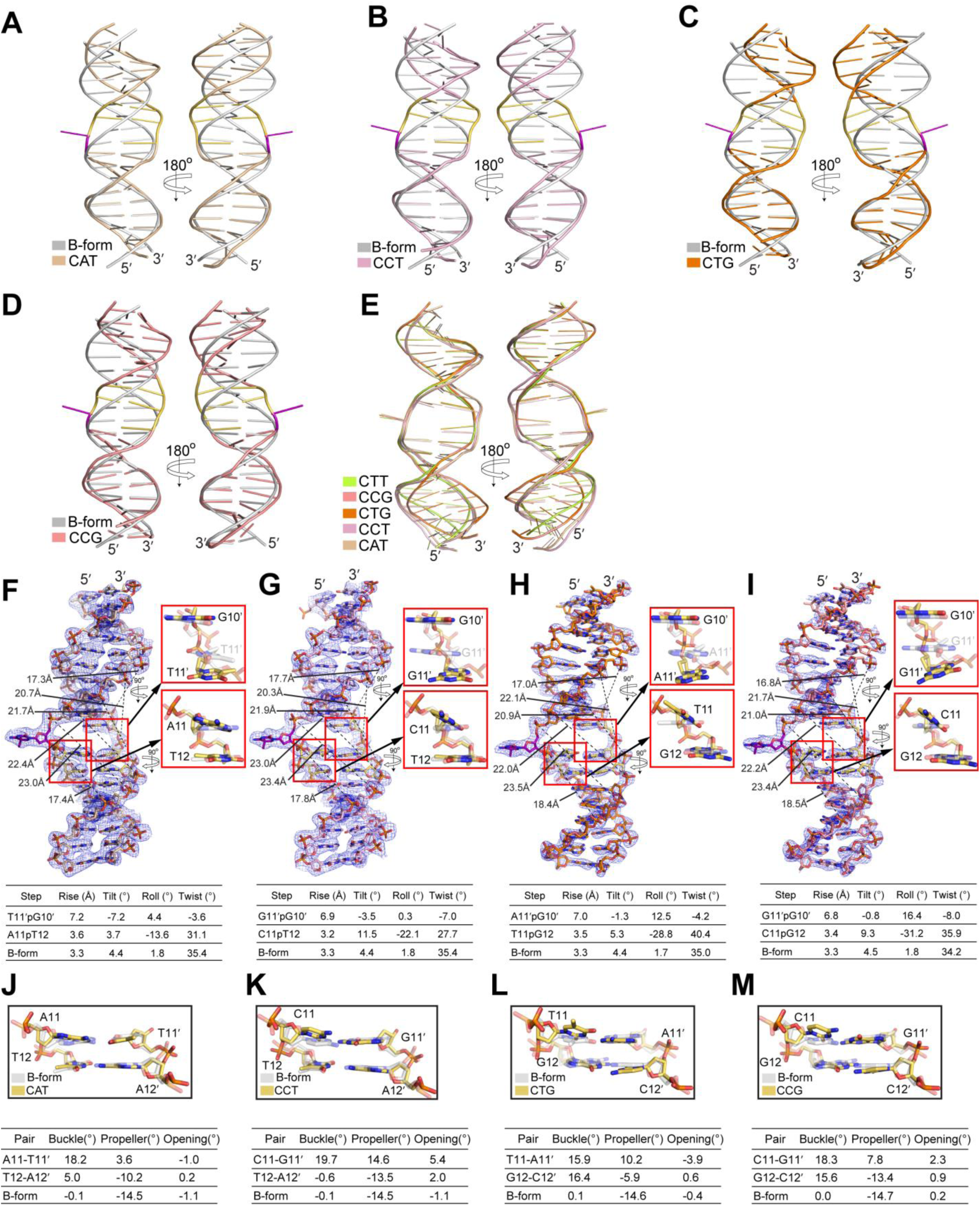
DRM2 binding-induced DNA deformation. (**A-D**) Structural superposition of the DRM2-bound CAT (A), CCT (B), CTG (C) and CCG (D) DNAs and the ideal B-form DNA with the corresponding sequences. (**E**) Structural overlay of the DRM2-bound CTT DNA with DRM2-bound CAT, CCT, CTG and CCG DNAs. (**F-I**) Fo-Fc omit maps (blue) of the CAT (F), CCT (G), CTG (H) and CCG (I) DNAs, contoured at the 2.0σ level. The major-groove widths of the deformed DNA upon binding of DRM2 are measured and labeled with dash lines. The base stacking between orphan Gua and +1 flanking nucleotide on the non-target strand and between the +1 and +2-flanking nucleotides on the target strand is shown in expanded views. The corresponding DNA base step parameters are listed below, compared to those for the ideal B-form DNA with corresponding sequences. (**J-M**) Structural overlay of the two base pairs next to fC10 between DRM2-bound CAT (J), CCT (K), CTG (L) or CCG (M) DNAs and the corresponding B-form DNA (gray). Detailed base-pair parameters are listed below, compared with those for corresponded ideal B-form DNA.

**Figure S5.**
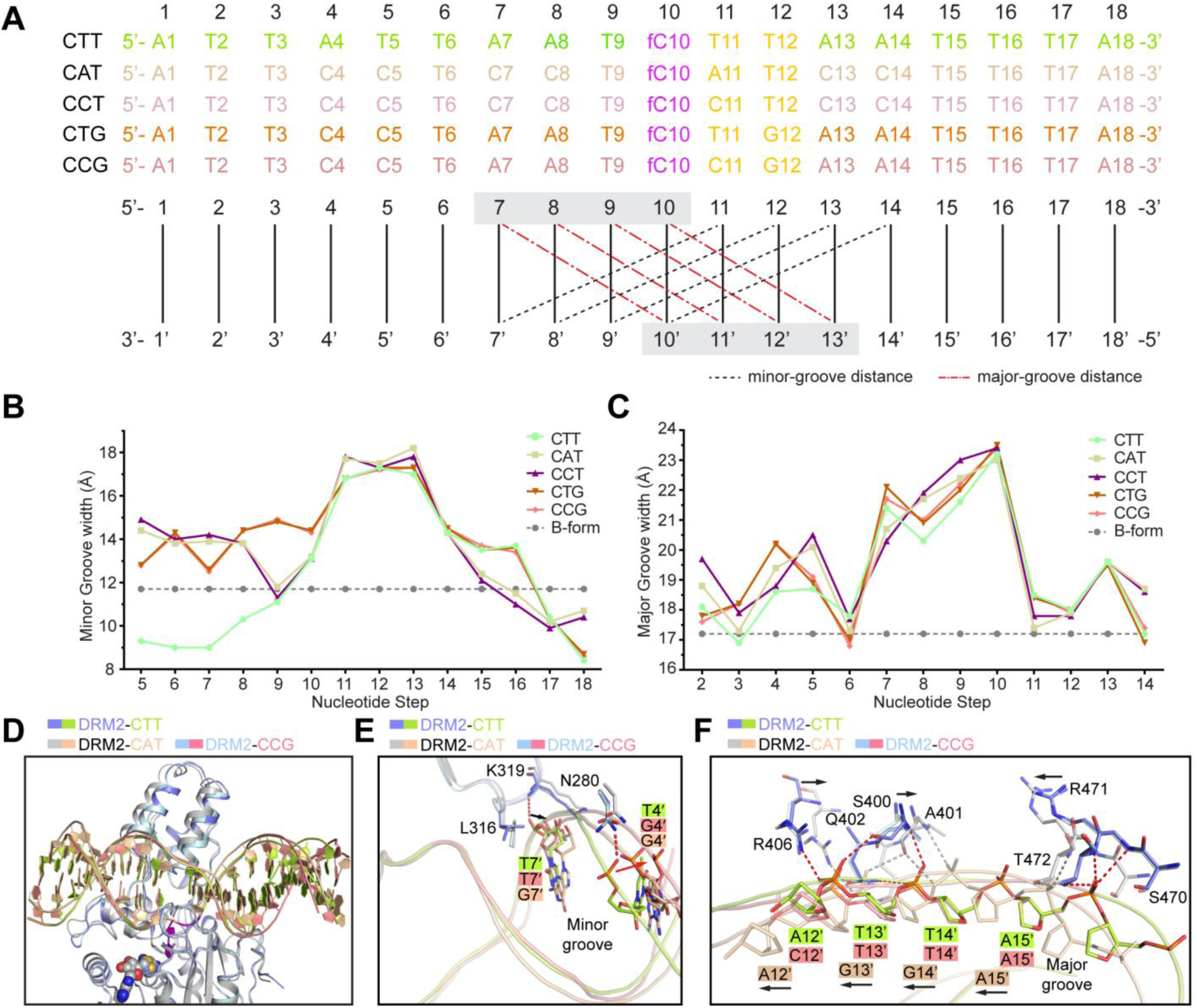
DNA shape analysis of DRM2-bound DNAs. (**A**) DNA sequences of the target strand for methylation in each structure. Minor groove widths (dashed lines in black) and major groove widths (dashed lines in red) are determined by measuring the cross-strand distances of the two phosphate groups of the nucleotides at indicated positions, as illustrated in the schematic below the DNA sequences. (**B-C**) DNA sequence-dependent minor groove widths (B) and major groove widths (C) of the DRM2-bound DNAs in the DRM2-CTT, DRM2-CAT, DRM2-CCT, DRM2-CTG and DRM2-CCG complexes. Dashed lines show the canonical groove widths for B-form DNA. (**D**) Structural overlay of the DRM2-CTT, DRM2-CAT and DRM2-CCG complexes highlight distinct shapes of the bounds DNAs. (**E**) Close-up view of the overlaid structures of the DRM2-CTT, DRM2-CAT and DRM2-CCG complexes, highlighting distinct minor-groove contacts of the N-terminal loop (N280) and the rearranged loop (L316 and K319) at the 5′ flanking region. (**F**) Close-up view of the overlaid structures of the DRM2-CTT, DRM2-CAT and DRM2-CCG complexes, highlighting distinct major-groove contacts of the LHH (S400-Q402 and R406) and the αH (S470-T472) at the 3′ flanking region.

**Figure S6.**
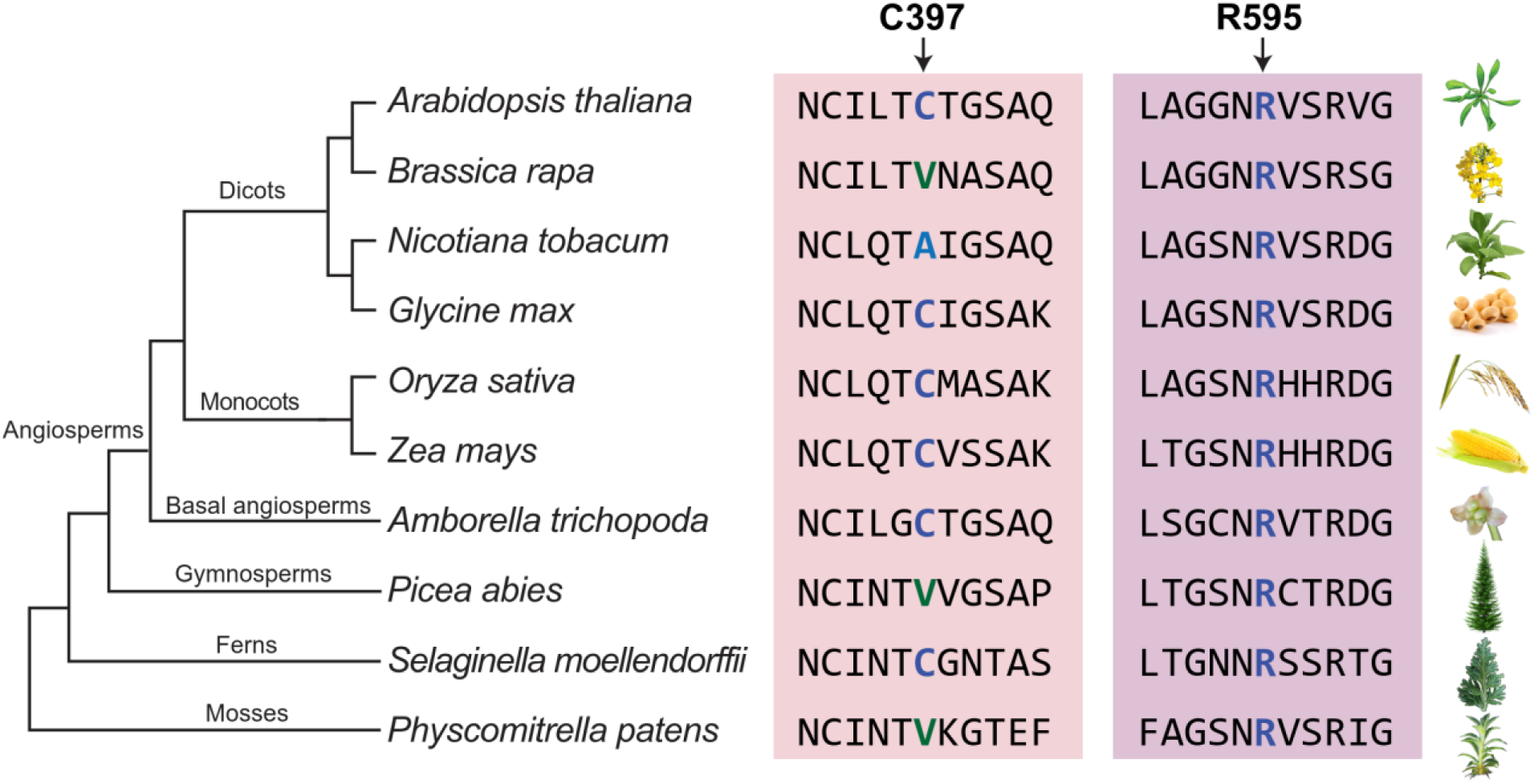
Phylogenetic tree showing conservation of C397 and R595 residues in DRM2 of different plant species. The amino acid sequences of DRM2 from plant species including mosses, ferns, gymnosperms and angiosperms are retrieved from Phytozyme (phytozome.jgi.doe.gov). The residues corresponding to C397 and R595 in DRM2 are highlighted.

**Figure S7.**
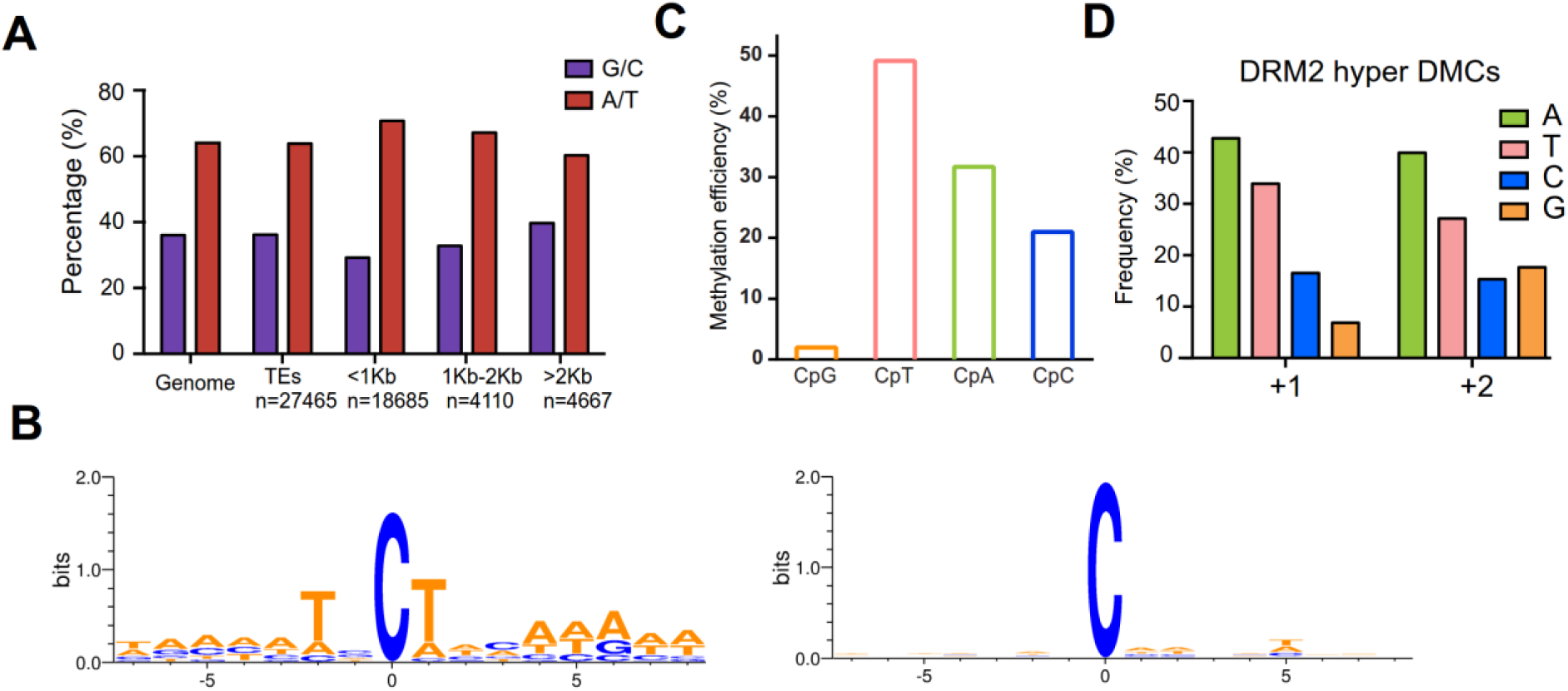
*In vitro* methylation analysis of the context-dependent DNA methylation by DRM2. (**A**) Histogram showing the percentage of A/T vs G/C composition of the Arabidopsis genome in comparison to all TEs and TEs of differing sizes. (**B**) Motifs of the top 21 over 237 total methylation sites on DNA fragments subjected to methylation by DRM2 and bisulfite sequencing analysis (left) or in native sequence (right). In total 28 DNA clones were used for sequencing analysis. (**C**) Summary of the methylation efficiencies of DRM2 on indicated CpG/CpH sites from a bisulfite-based methylation assay using a 637-bp DNA fragment, biological replicate of Fig. 2I. In total 8 clones were sequenced for analysis. (**D**) Quantification of the nucleotide frequency of the first nucleotide downstream (+1) and the second nucleotide downstream (+2) of hyper DRM2 DMCs called against *ddc*.

**Figure S8.**
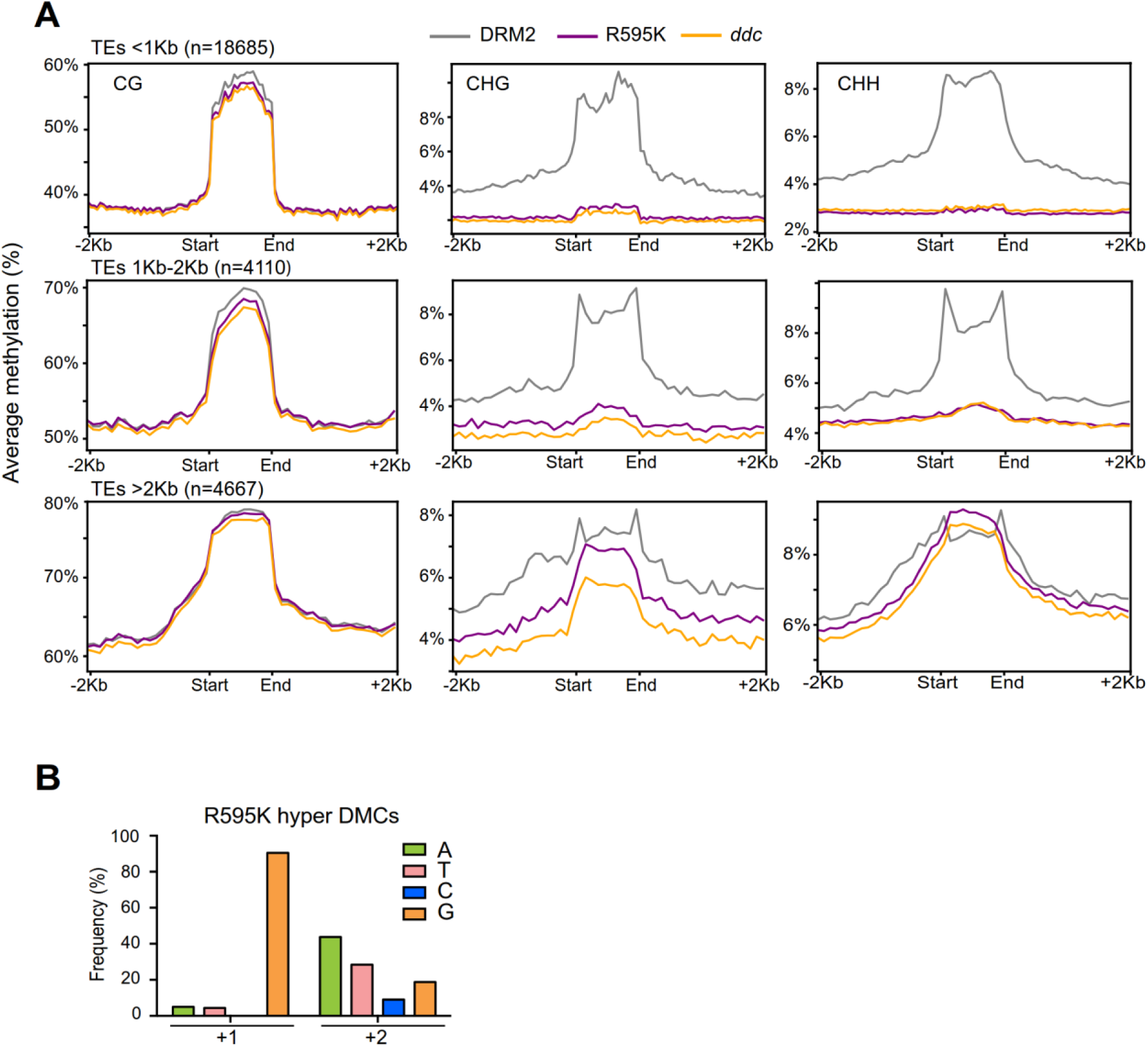
Genomic methylation analysis of DRM2 and R595K. (**A**) Average CG, CHG, and CHH methylation levels of DRM2 and R595K over transposable elements (TEs) of different sizes. (**B**) Quantification of the nucleotide frequency of the first nucleotide downstream (+1) and the second nucleotide downstream (+2) of all hyper R595 DMCs called against *ddc*.

**Figure S9.**
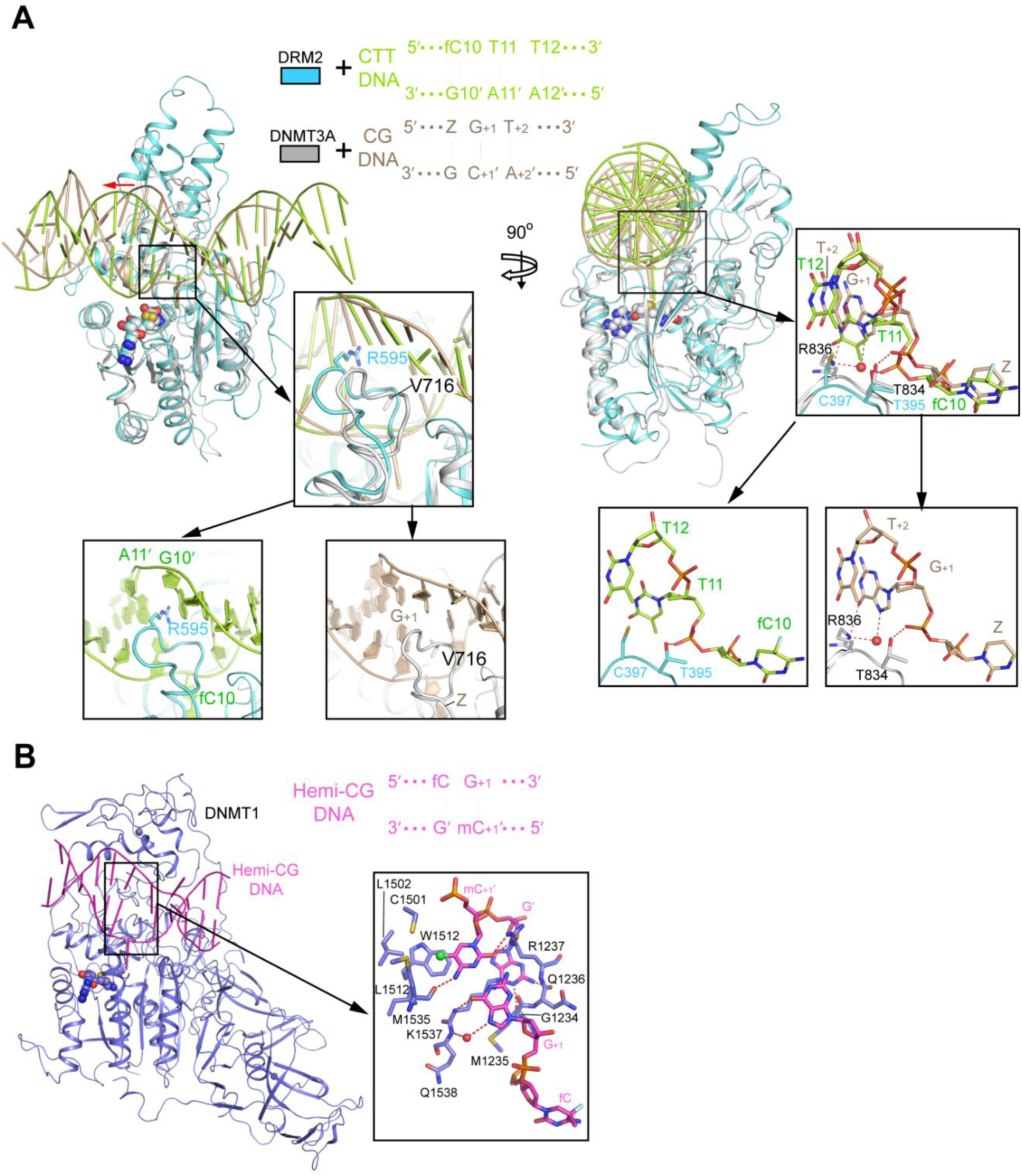
Structural comparison of enzyme-substrate complexes of DRM2 and mammalian DNA methyltransferases. (**A**) Structural overlay of DRM2-CTT complex (cyan) and DNMT3A-CG DNA complex (gray, PDB: 5YX2), with the expanded views highlighting the DNA interactions of the catalytic loop (left) and the TRD loop (right). Note that the DRM2-DNA complex contains a 25-mer DNA duplex spanning two DNMT3A molecules. For clarity, only one DNMT3A molecule and associated DNA are shown. Z, Zebularine, a cytosine analogue. (**B**) Structure of mouse DNMT1 (731-1602) bound to hemi-methylated CG DNA (PDB: 4DA4), with the DNA-interaction site on the catalytic loop shown in expanded views.

**Figure S10.**
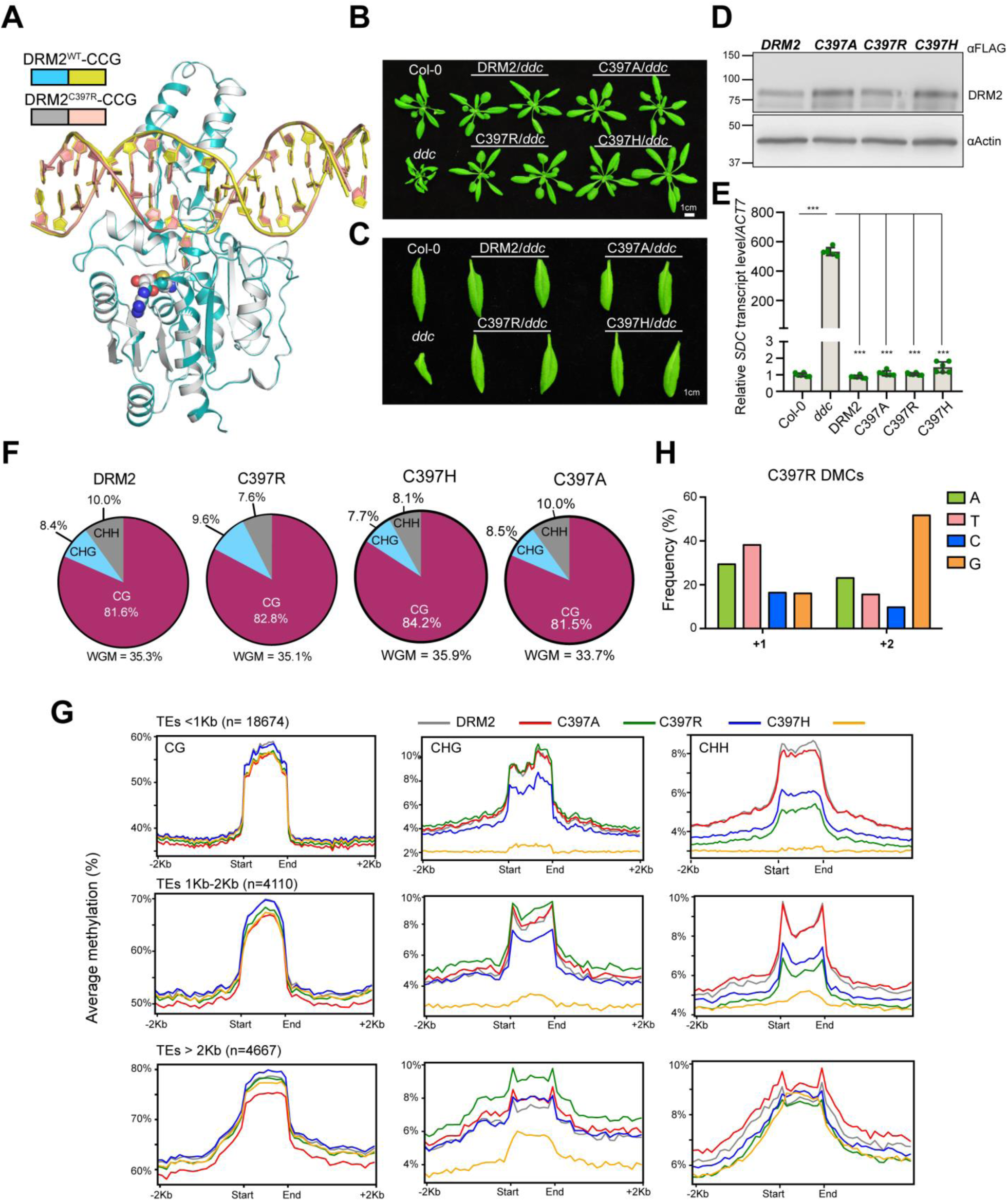
The C397R mutation shifts the substrate specificity of DRM2 toward CHG DNA. (**A**) Structural overlay of WT (gray) and C397R (cyan)-mutated DRM2 in complex with CCG DNA. The bound DNAs are colored in yellow and wheat, respectively. The SAH molecules are shown in sphere representation. (**B**) Phenotypes of 3-week-old plants. Col-0 and *ddc* (*drm1drm2cmt3*) serve as controls. (**C**) Single leaf phenotypes of lines shown in panel (A). (**D**) Western blot of FLAG-tagged DRM2 proteins from same lines listed in panel (A). (**E**) RT-qPCR of *SDC* relative transcript level. Data are mean ± SD. Statistical analysis used two-tailed Student’s t-test for the difference from WT. ***, p < 0.001. (**F**) Pie charts showing the genome wide proportion of each methylation type in DRM2, C397R, C397A and C397H. (**G**) Metaplots showing average CG, CHG, and CHH methylation for DRM2, C397A, C397R, C397H, and *ddc* over transposable elements (TEs) of different sizes. (**H**) Quantification of the frequency (%) of each nucleotide in the +1 base pair and +2 base pair downstream of C397R hyper DMCs depicted in Fig. 4F.

**Table S1.** X-ray data collection and refinement statistics.

|  | DRM2-CTT<br>DNA<br>(PDB: xxxx) | DRM2-CAT<br>DNA<br>(PDB: xxxx) | DRM2-CCT<br>DNA<br>(PDB: xxxx) | DRM2-CTG<br>DNA<br>(PDB: xxxx) | DRM2-CCG<br>DNA<br>(PDB: xxxx) | DRM2<sup>C397R</sup>-<br>CCG DNA<br>(PDB: xxxx) |
| --- | --- | --- | --- | --- | --- | --- |
| **Data collection** |  |  |  |  |  |  |
| **Space group** | C 2 2 21 | C 1 2 1 | C 1 2 1 | C 2 2 21 | C 2 2 21 | C 2 2 21 |
| **Cell dimensions** |  |  |  |  |  |  |
| *a, b, c* (Å) | 54.4, 232.5,<br>117.8 | 172.2, 70.8,<br>103.7 | 171.9, 71.0,<br>104.1 | 116.9, 230.4,<br>54.2 | 116.9, 230.4,<br>54.2 | 118.3 231.7,<br>54.3 |
| *α, β, γ* (°) | 90, 90, 90 | 90, 98.0, 90 | 90, 97.9, 120 | 90, 90, 90 | 90, 90, 90 | 90, 90, 90 |
| **Resolution** (Å) | 48.29-2.11<br>(2.17-2.11) | 102.7-2.55<br>(2.65-2.55) | 44.33-2.80<br>(2.90-2.80) | 115.2-2.56<br>(2.68-2.56) | 49.2-2.61<br>(2.73-2.61) | 49.2 - 2.25<br>(2.33-2.25) |
| *R*<sub>merge</sub> | 0.119 (1.814) | 0.08 (1.385) | 0.098 (1.0) | 0.139 (1.198) | 0.17 (1.439) | 0.155 (1.625) |
| *I*/*σ*(*I*) | 10.7 (0.9) | 14.0 (1.0) | 6.4 (1.0) | 7.3 (1.2) | 7.6 (0.8) | 7.8 (0.8) |
| *CC*<sub>1/2</sub> | 0.997 (0.337) | 0.999 (0.413) | 0.993 (0.536) | 0.993 (0.592) | 0.990 (0.409) | 0.966 (0.444) |
| **Completeness** (%) | 99.2 (98.1) | 99.7 (99.9) | 99.0 (94.8) | 98.6 (94.3) | 99.5 (99.3) | 97.02 (93.16) |
| **Redundancy** | 5.8 (5.3) | 6.7 (6.7) | 3.5 (3.3) | 4.4 (4.4) | 5.0 (4.3) | 7.2 (6.8) |
| **Refinement** |  |  |  |  |  |  |
| **No. reflections** | 43158 | 40429 | 30434 | 23628 | 22956 | 35099 |
| *R*<sub>work</sub> / *R*<sub>free</sub> | 0.196/0.220 | 0.225/0.263 | 0.214/0.263 | 0.211/0.249 | 0.213/0.240 | 0.180/0.210 |
| **No. atoms** |  |  |  |  |  |  |
| **Protein/DNA** | 3548 | 6911 | 6848 | 3549 | 3549 | 3554 |
| **Ligand** | 26 | 52 | 52 | 26 | 26 | 26 |
| **Water** | 378 | 54 | 12 | 126 | 57 | 343 |
| ***B* factors (Å<sup>2</sup>)** |  |  |  |  |  |  |
| **Protein** | 42.6 | 103.2 | 95.7 | 60.5 | 53.2 | 41.9 |
| **DNA** | 61.5 | 141.3 | 145.2 | 83.9 | 80.7 | 66.9 |
| **Ligand** | 33.7 | 82.5 | 75.5 | 51.0 | 43.7 | 33.8 |
| **Water** | 48.9 | 85.3 | 61.2 | 56.9 | 49.4 | 49.0 |
| **r.m.s deviations** |  |  |  |  |  |  |
| **Bond lengths** (Å) | 0.004 | 0.004 | 0.004 | 0.004 | 0.003 | 0.005 |
| **Bond angles** (°) | 0.84 | 0.64 | 0.63 | 0.62 | 0.60 | 0.76 |

<sup>a</sup>Values in parentheses are for highest-resolution shell. Each structure was determined using the dataset collected from a single crystal.

**Table S2.** Bisulfite-sequencing statistics data.

\* Fold coverage was calculated by multiplying the unique reads by the read length
| Sample | Total # reads | Aligned reads | % alignment | Unique reads | % unique | Covered cytosines | Fold coverage* | Bisulfite Conversion (%) |
| --- | --- | --- | --- | --- | --- | --- | --- | --- |
| DRM2 | 92254114 | 81855508 | 89 | 66560025 | 72 | 40045047 | 28 | 99.6 |
| C397A | 101296272 | 52868014 | 52 | 43378390 | 43 | 38504208 | 18 | 99.6 |
| C397R | 80912583 | 78329769 | 97 | 64868492 | 80 | 39311904 | 27 | 99.7 |
| C397H | 69913065 | 68478853 | 98 | 57067847 | 82 | 4098617 | 24 | 99.4 |
| R595K | 72563481 | 70780428 | 98 | 58547677 | 81 | 42451767 | 25 | 99.5 |
| *ddc* | 86549847 | 84121668 | 97 | 70258321 | 81 | 40638219 | 30 | 99.4 |

(50bp) and dividing it by the size of the Arabidopsis genome (119Mb).

**Table S3.** Number of hyper differentially methylated cytosines (DMCs).

| Sample | CG DMC | CHG DMC | CHH DMC |
| --- | --- | --- | --- |
| DRM2/*ddc* | 6792 | 16087 | 76234 |
| C397R/*ddc* | 4722 | 14002 | 10623 |
| R595K/*ddc* | 1180 | 18 | 107 |

